# Two new hybrid zones expand the swordtail hybridization model system

**DOI:** 10.1101/2020.11.18.389205

**Authors:** Daniel L. Powell, Ben Moran, Bernard Kim, Shreya M. Banerjee, Stepfanie M. Aguillon, Paola Fascinetto-Zago, Quinn Langdon, Molly Schumer

## Abstract

Natural hybridization events provide unique windows into the barriers that keep species apart as well as the consequences of their breakdown. Here we characterize hybrid populations formed between the northern swordtail fish *Xiphophorus cortezi* and *X. birchmanni* from collection sites on two rivers. We develop sensitive and accurate local ancestry calling for this system based on low coverage whole genome sequencing. Strikingly, we find that hybrid populations on both rivers consist of two genetically distinct subpopulations: a cluster of nearly pure *X. birchmanni* individuals and one of phenotypically intermediate hybrids that derive ~85-90% of their genome from *X. cortezi*. Simulations and empirical data suggest that at both sites initial hybridization occurred ~150 generations ago, with little evidence for contemporary gene flow between subpopulations, likely due to strong assortative mating. The patterns of population structure uncovered here mirror those seen in hybridization between *X. birchmanni* and its sister species, *X. malinche*. Future comparisons will provide a window into the repeatability of the outcomes of hybridization not only across independent hybridization events between the same species but also across distinct species pairs.

## Introduction

It has long been recognized that hybrids provide unique insights into the barriers between species and the consequences of their breakdown (Barton & Hewitt, 1985). While artificial hybrids, particularly in *Drosophila*, formed the foundation of early research into the genetic barriers that differentiate species (Coyne & Orr, 1997; Dobzhansky, 1936; Orr & Coyne, 1989), in recent years there has been a renaissance in the study of natural hybrid populations (e.g. Brandvain, Kenney, Flagel, Coop, & Sweigart, 2014; Powell et al., 2020; Sankararaman et al., 2014; Stukenbrock, Christiansen, Hansen, Dutheil, & Schierup, 2012; Turissini & Matute, 2017). These natural experiments provide the unique opportunity to study hybridization in its ecological and evolutionary contexts, which are fundamental to fully characterizing consequences of hybridization (Barton & Hewitt, 1985).

More recently, the increasing accessibility of dense genomic data has allowed granular studies of genome evolution in hybrid zones, revealing variation in ancestry among individuals and populations as well as selection on ancestry at particular loci (Taylor, Larson, & Harrison, 2015; Teeter et al., 2008; Torre, Ingvarsson, & Aitken, 2015). This increased resolution has allowed researchers to begin to compare distinct hybridization events, a first step towards tackling the important question of how repeatable outcomes of hybridization are at both the population and genomic level.

Though the complexity of natural hybrid zones provides an opportunity to study the interactions of different genetic, ecological, and evolutionary forces, it also creates challenges in disentangling them. For example, it can be difficult to determine whether patterns observed in individual hybrid zones are driven by intrinsic interactions between the genomes of hybridizing species, dependent upon demographic and ecological context, or are stochastic (Ross & Harrison, 2002). Thus, the study of independent hybrid zones provides the best of both worlds, with natural replication testing the repeatability of evolution after hybridization, and variation in environment or demographic history between populations creating informal tests for the relevance of these factors (Harrison & Larson, 2016; Janoušek et al., 2012). Each new case described offers a unique window into how eco-evolutionary history drives hybridization outcomes.

Due in part to their natural replication in multiple river systems, hybrid populations formed between swordtail fish *Xiphophorus birchmanni* and *X. malinche* have become an emerging model for the study of hybridization (Rosenthal et al., 2003). Research in this system has revealed that hybridization between *X. birchmanni* and *X. malinche* began recently in several populations (Schumer et al., 2014), likely due to disrupted sensory communication as a result of human-mediated habitat disturbance (Fisher, Wong, & Rosenthal, 2006). Moreover, our work has indicated that differences in the strength of assortative mating by ancestry explain differences in population structure between *X. birchmanni* x *X. malinche* hybrid populations in distinct rivers (Schumer et al., 2017).

Here, we describe a previously unexplored hybridization event between *X. birchmanni* and its more distant relative, *X. cortezi* (Kallman & Kazianis, 2006). We characterize the history of hybridization in two geographically independent tributaries of the Río Santa Cruz drainage in northern Hidalgo, Mexico. Using sensitive and accurate local ancestry calling, we infer demographic history of each population and evaluate the role of assortative mating in maintaining ancestry structure. Like *X. birchmanni* and *X. malinche*, the two species have overlapping ranges but largely are separated along an elevational gradient. *X. malinche* occurs at the highest elevations of all three species. In streams where *X. birchmanni* and *X. cortezi* co-occur, *X. cortezi* is found at lower elevations. Moreover, both pairs of hybridization events include a sworded (*X. malinche*; *X. cortezi*) and swordless species (*X. birchmanni*), among other differences in sexual signals (Cui, Delclos, Schumer, & Rosenthal, 2017; Culumber & Rosenthal, 2013; Fernandez & Morris, 2008; Rosenthal et al., 2003). These analyses allow us to characterize cases of hybridization which are independent in their evolutionary history, yet broadly parallel in the recent onset and ongoing nature of hybridization, providing a valuable opportunity to study the population-level outcomes of hybridization in closely related species pairs. As such, these new hybrid populations provide a powerful window into the barriers between species and the consequences of their breakdown.

## Materials and Methods

### Sample collection

Fish were collected from wild populations in the states of Hidalgo and San Luis Potosí, Mexico using baited minnow traps. Putative *X. cortezi* x *X. birchmanni* hybrids were sampled from two distinct collection sites (hereafter sites; Fig. 1), Huextetitla (21°9’43.82”N 98°33’27.19”W, n=87) and Santa Cruz (21°9’27.63”N 98°31’13.79”W, n=95). These sites occur in separate tributaries of the Río Santa Cruz in northern Hidalgo. Pure *X. cortezi* were collected in January 2020 from the Río Huichihuayán, a fully allopatric population with respect to *X. birchmanni* (Puente de Huichihuayán, 21°26’9.95”N 98°56’0.00”W, n=42). One previously sequenced pure *X*. *cortezi* individual from Las Conchas (21°23’33.30”N 98°59’23.33”W) and seven from el nacimiento de Huichihuayán (21°27’34.10”N 98°58’36.70”W) were included in analyses (Powell et al., 2020; Schumer et al., 2018). Likewise, pure *X. birchmanni* from Coacuilco (21° 5’51.16”N 98°35’20.10”W) were collected previously for studies of hybridization with *X. malinche* (Schumer et al., 2018).

**Figure 1.**
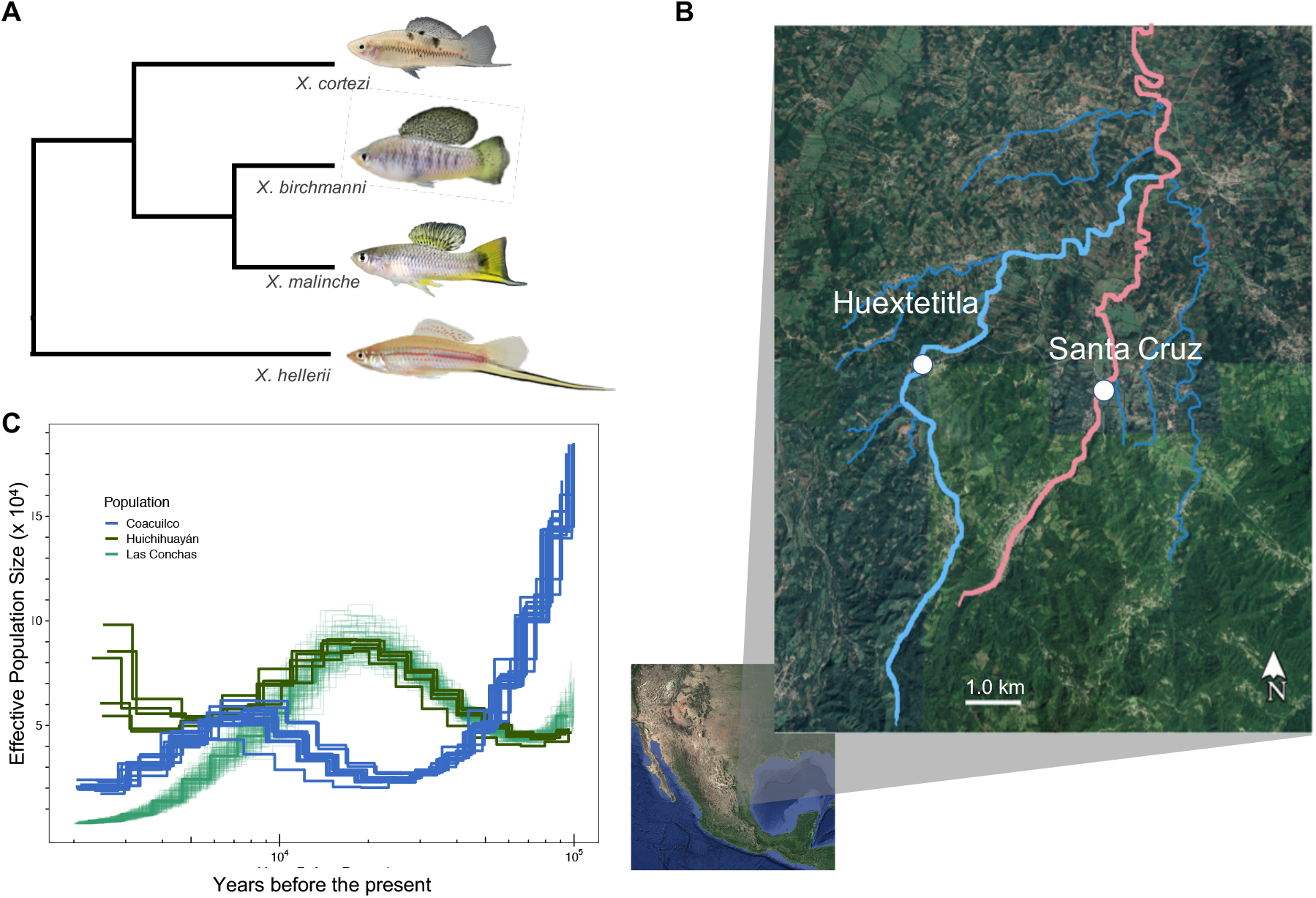
Phylogenetic relationships between species, location of sampling sites, and demographic history of parental species. **A**) Phylogenetic relationships between *X. birchmanni*, *X. malinche*, and *X. cortezi* (simplified from Cui et al., 2013). **B**) Sampling sites for *X. birchmanni* x *X. cortezi* hybrid populations with the tributaries they occur in highlighted. **C**) Demographic history of pure *X. birchmanni* (blue lines) and *X. cortezi* (green lines) populations inferred by PSMC, assuming 2 generations a year and a per-base pair mutation rate of 3.5 × 10^−9^.

After collection, fish were anesthetized in a buffered solution of MS-222 and water diluted to 100 mg/mL (Stanford APLAC protocol #33071). Once anesthetized fish were photographed against a grid background with dorsal and caudal fins spread using a Nikon d90 DSLR digital camera mounted to a copy stand and equipped with a macro lens. A small fin clip was taken from each individual and preserved in 95% ethanol for later DNA extraction. Females used for embryo comparisons were euthanized by MS-222 overdose in the field before being preserved in 95% ethanol.

### Phenotyping and PCA analysis

Standard length, body depth, peduncle depth, caudal fin length, dorsal fin width, dorsal fin height were measured from photographs of adult fish using ImageJ (Schneider, Rasband, & Eliceiri, 2012). Additionally, length of the sword, a sexually selected male ornament that differs between *X. birchmanni* and *X. cortezi*, was measured for adult males. Principal Component Analysis (PCA) was performed on males and females separately to compare phenotypic variance in the hybrid populations to phenotypic variance in the parental species using the *princomp* function in R v.3.6.3. Hybrid individuals were assigned as part of the *X. birchmanni* or *X. cortezi* ancestry clusters based on their genome-wide ancestry proportions (see *Inferring Local Ancestry Section* below).

Discriminant function analysis was performed to assess how well morphological phenotypes could predict ancestry in males. A linear discriminant analysis model was trained using 75% of individuals categorized based on their genome-wide ancestry as pure *X. birchmanni,* pure *X. cortezi, X. birchmanni*-like hybrid or *X. cortezi*-like hybrid, and was then used to predict the ancestry category of the remaining 25% of individuals. Training on different subsets ranging from 30-90% did not qualitatively change results.

### DNA extraction and library preparation

DNA was extracted from fin tissue using the Agencourt DNAdvance kit (Beckman Coulter, Brea, California) as specified by the manufacturer but using half the recommended reaction volume. Extracted DNA was quantified using the TECAN Infinite M1000 microplate reader (Tecan Trading AG, Switzerland) at the High Throughput Biosciences Center at Stanford University, Stanford, CA.

Tagmentation-based whole genome libraries for low coverage sequencing were prepared from DNA extracted from fin clips collected from fish caught at the Huextetitla and Santa Cruz populations. Briefly, DNA was diluted to approximately 2.5 ng/μL and enzymatically sheared using the Illumina Tagment DNA TDE1 Enzyme and Buffer Kits (Illumina, San Diego, CA) at 55°C for 5 minutes. Sheared DNA samples were amplified in PCR reactions with dual indexed custom primers for 12 cycles. Amplified PCR reactions were pooled and purified using 18% SPRI magnetic beads.

Genomic libraries for high coverage sequencing of an individual collected in Huextetitla was prepared following Quail et al. (Quail, Swerdlow, & Turner, 2009). Briefly, approximately 500 ng of DNA was sheared to ~400 basepairs using a QSonica sonicator (QSonica Sonicators, Newton, Connecticut). To repair the sheared ends, DNA was mixed with dNTPs and T4 DNA polymerase, Klenow DNA polymerase and T4 PNK and incubated at room temperature for 30 minutes (NEB, Ipswich, MA) and then purified with the Qiagen QIAquick PCR purification kit (Qiagen, Valencia, CA). A-tails were added by mixing the purified end-repaired DNA with dATPs and Klenow exonuclease and incubating at 37° C for 30 minutes (NEB, Ipswich, MA) and then purified using the Qiagen QIAquick PCR purification kit (Qiagen, Valencia, CA). Adapter ligation reaction was performed followed by purification with the Qiagen QIAquick PCR purification kit (Qiagen, Valencia, CA). Adapter ligated DNA was amplified using indexed primers in individual Phusion PCR reactions for 12 cycles and then purified using 18% SPRI beads.

Libraries were quantified with a Qubit fluorometer (Thermo Scientific, Wilmington, DE). Library size distribution and quality were assessed using Agilent 4200 Tapestation (Agilent, Santa Clara, CA). Libraries were sequenced on an Illumina HiSeq 4000 at Admera Health Services, South Plainfield, NJ.

### 10X chromium library

To generate a draft assembly for *X. cortezi*, we made a 10X Chromium library using the Genomic Services Lab at the HudsonAlpha Institute for Biotechnology. High molecular weight DNA was extracted from fin tissue using the Genome Reagent Kit from 10X genomics. DNA was diluted to working concentrations of 0.4 ng/μL, quantified with a Qubit fluorometer. This is the recommended concentration given the *Xiphophorus* genome size of ~700 Mb. These working solutions were used as input to the library preparation protocol to begin the emulsion phase. The emulsion phase was broken as directed by the protocol, and bead purification was performed in 96-well plates. Final libraries were quantified using a Qubit fluorometer and library size was evaluated on a Bioanalyzer.

### Admixtools analysis to evaluate evidence for hybridization between X. birchmanni and X. cortezi

To evaluate initial evidence for admixture, we sequenced one individual from Huextetitla who appeared phenotypically intermediate between *X. birchmanni* and *X. cortezi* to ~30X coverage, as described above. We mapped reads from this individual to the *X. birchmanni* reference genome using bwa (Li & Durbin, 2009), marked and removed duplicates with Picard Tools and realigned insertion-deletion differences (indels) with GATK v3.4 (McKenna et al., 2010). We performed variant calling with GATK’s HaplotypeCaller in GVCF mode (McKenna et al., 2010). Because we lack an appropriate variant set for variant recalibration, we did not perform this step and instead implemented hard-calls based on several filters (DP, QD, MQ, FS, SOR, ReadPosRankSum, and MQRankSum) as described previously (Schumer et al., 2018). In addition, we masked 5 bp windows surrounding indels and any site with greater than 2X or less than 0.5X the average genome-wide coverage. Based on past work quantifying Mendelian errors in swordtail pedigrees after applying these filters, we believe that this approach has high accuracy (Schumer et al., 2018).

We repeated these steps for previously sequenced *X. malinche*, *X. birchmanni*, and *X. cortezi* individuals to generate variant calls from an appropriate set of species for D-statistic analysis (Patterson et al., 2012). We used custom scripts available on our lab github to convert these files to admixtools format (https://github.com/Schumerlab/Lab_shared_scripts; https://openwetware.org/wiki/Schumer_lab:_Commonly_used_workflows#g.vcf_files_to_Admixtools_input). This resulted in 1,001,493 informative sites for analysis with admixtools. We used the qpDstat function from admixtools and a jack-knife bootstrap window size of 5 Mb to determine the most likely four-population tree, and calculate the D-statistic based on that tree. We also explored evidence of admixture with another *Xiphophorus* species that is sympatric with *X. birchmanni* and *X. cortezi* but deeply diverged from both species and found no evidence for hybridization with this species (Supporting Information 1-2).

### Generation of a reference guided X. cortezi assembly

An initial draft assembly for *X. cortezi* was generated from the 10X Chromium library described above using the supernova software (v2.0.1; Weisenfeld, Kumar, Shah, Church, & Jaffe, 2017). The maximum reads used parameter was set to 280 million and the output style was specified as pseudohap, otherwise recommended parameters for the *Xiphophorus* genome size were used. This resulted in a draft assembly of 7,610 scaffolds (2,182 longer than 10 kb) with an N50 of 1.04 Mb and a total of 686 Mb assembled. The expected genome size of *Xiphophorus* is approximately 700 Mb.

Chromosome-scale synteny is conserved as 24 chromosomes across *Xiphophorus* species (Amores et al., 2014; Powell et al., 2020; Schartl et al., 2013). Thus, we decided to leverage the chromosome structure in other *Xiphophorus* assemblies to create chromosome-level scaffolds for *X. cortezi*. First, we created a multi-way whole genome alignment for swordtail species including *X. birchmanni, X. variatus*, and *X. malinche* (Powell et al., 2020), *X. cortezi* and *X. xiphidium* (this study), *X. couchianus* (RefSeq assembly GCF_001444195.1), and *X. maculatus* (RefSeq assembly GCF_002775205.1). Using the phylogenetic relationships from Cui et al. (2013) as our guide tree, we ran progressive Cactus (Armstrong et al., 2019) to build the alignment. Parameters for the alignment are automatically determined by progressive Cactus based on branch lengths of the guide tree. Using this alignment and the same guide tree described previously, we arranged the scaffolds into 24 putative chromosomes using Ragout (Komolgorov et al. 2018), keeping the naming scheme consistent with that of the *X. birchmanni* genome (Fig. S1). Chromosome aligned scaffolds (N=28) were combined with unplaced scaffolds (N=4,777) to create the final assembly. Configuration files and associated scripts, as well as a Docker environment, are provided on github at https://github.com/Schumerlab/Xbir_xcor_hybridzone.

### PSMC demographic inference

We inferred the demographic history of *X. cortezi* using the 10X Chromium library generated for the *X. cortezi* genome assembly, as well as previously sequenced *X. cortezi* individuals from Huichihuayán and *X. birchmanni* individuals from Coacuilco (Powell et al., 2020; Schumer et al., 2018). Briefly, raw reads were mapped to the *X. birchmanni* reference assembly (Powell et al., 2020), after which GATK v3.4 (McKenna et al., 2010) was used to call variant sites as described above. These variants were then quality filtered as described above and used to create pseudo-reference genomes for each individual, which were input to PSMC (Li & Durbin, 2011). PSMC output was converted to effective population size assuming a mutation rate of 3.5×10^−9^ bp^−1^ generation^−1^ and a generation time of 0.5 years, as described previously (Schumer et al., 2018). We note that although other methods such as MSMC allow for simultaneous inference of demographic history in multiple individuals, they also require phasing, which can introduce errors, especially in cases where high quality reference panels are not available (Schiffels & Wang, 2020).

### Inferring local ancestry

We used a series of approaches to develop ancestry informative sites that distinguished *X. birchmanni* and *X. cortezi*. We first used a panel of 25 high coverage *X. birchmanni* individuals from the Coacuilco population, 7 *X. cortezi* individuals from el nacimiento de Huichihuayán, and the reference individual from Las Conchas that were collected in previous work (Powell et al., 2020; Schumer et al., 2018) to identify candidate ancestry informative sites. With this candidate set and low coverage whole-genome sequence data that we collected for *X. cortezi* in this study (N=30) and previously had collected for *X. birchmanni* (Schumer et al., 2018), we evaluated population level counts for *X. cortezi* and *X. birchmanni* alleles at these ancestry informative sites. Any candidate ancestry informative site where the major allele in either parental population was at less than 90% frequency was excluded, yielding a set of 1.1 million ancestry informative sites genome-wide (~1.5 per kb). We describe our approach for identifying ancestry informative sites and determining parameters for local ancestry inference in more detail in Supporting Information 3-4; we have also explored these issues in previous work (Powell et al., 2020; Schumer, Powell, & Corbett-Detig, 2020).

With this set of ancestry informative sites, we used a hidden Markov model (HMM) approach to infer local ancestry with our previously developed local ancestry inference tool, *ancestryinfer* (Schumer et al., 2020), and evaluated performance on a set of parental individuals that were not used in previous steps (Fig. S2). We also performed simulations to evaluate expected performance under a range of demographic scenarios. Together these results suggest that we expect to have high accuracy in calling local ancestry in *X. birchmanni* x *X. cortezi* hybrids (Supporting Information 3-4; Fig. 2; Fig. S3).

**Figure 2.**
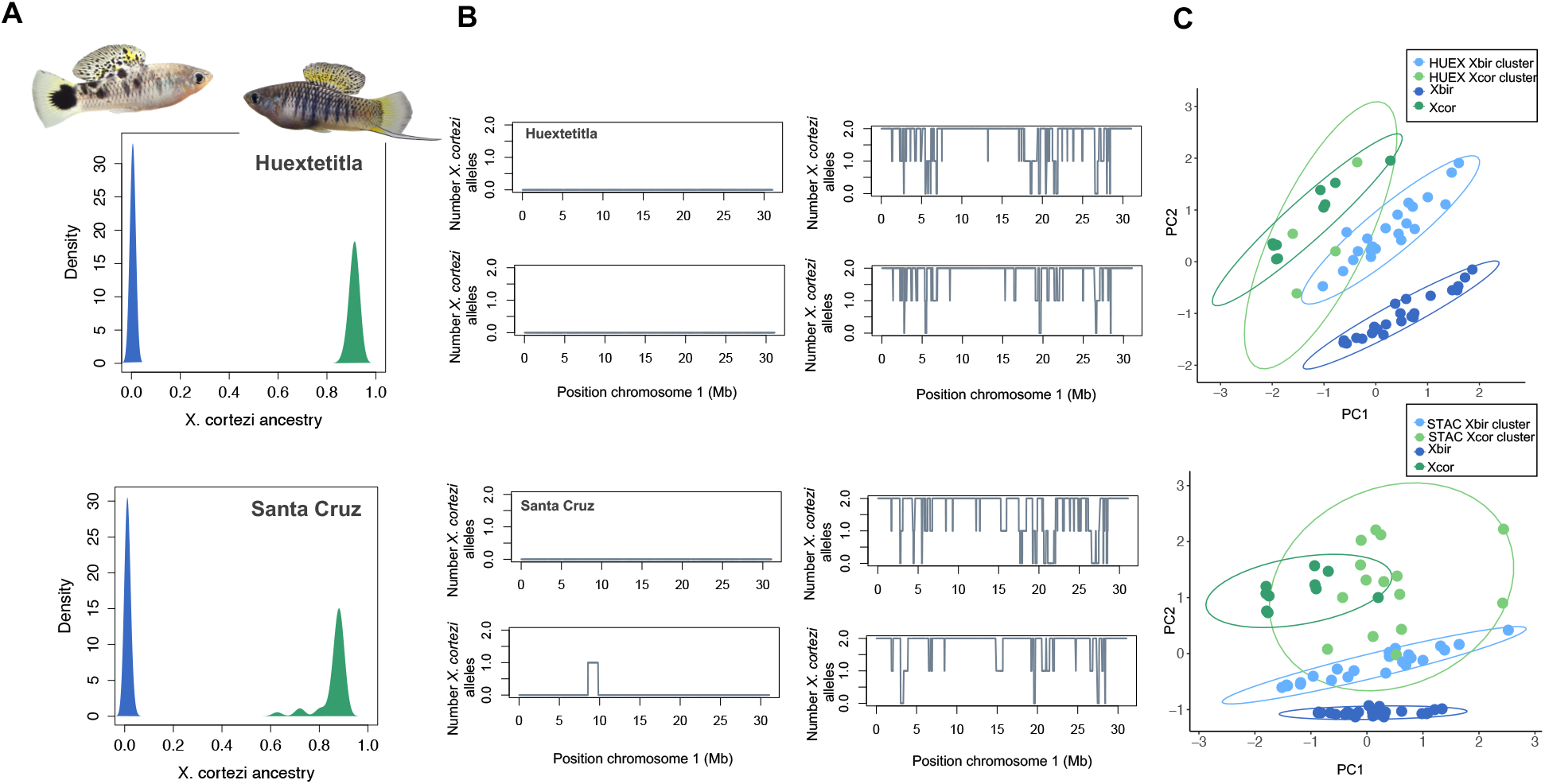
Ancestry structure of *X. birchmanni* x *X. cortezi* populations and example local ancestry inference. **A**) Genome-wide ancestry in the Huextetitla (top) and Santa Cruz (bottom) populations. Plotted here is the proportion of the genome derived from *X. cortezi* in all sampled individuals in the population. Individuals plotted in green were assigned to the *X. cortezi* ancestry cluster and in blue were assigned to the *X. birchmanni* ancestry cluster. Representative individuals from each ancestry cluster from the Huextetitla population are shown. **B**) Local ancestry on chromosome 1 for two *X. birchmanni* cluster (left) and *X. cortezi* cluster (right) individuals for the Huextetitla (top) and Santa Cruz (bottom) populations. **C**) PCA plots of phenotypic data from Huextetitla population males (top) and Santa Cruz population males (bottom) compared with parental species male phenotypic data. Xbir – *X. birchmanni*, Xcor – *X. cortezi*, HUEX – Huextetitla, STAC – Santa Cruz. Each point represents one individual and ellipses represent the 95% confidence interval. Loadings for each phenotype can be found in Table S1.

Confident in the accuracy of our local ancestry inference approach, we next applied these methods to individuals collected from putative hybrid populations. Based on the results of an initial analysis with uniform ancestry priors, we identified the presence of two distinct ancestry clusters at both collection sites (Supporting Information 3-4). We thus re-ran the HMM for each genetic cluster using cluster-specific ancestry priors (*X. birchmanni* cluster: 1% *X. cortezi; X. cortezi* hybrid cluster: 15% *X. birchmanni*) and generated a merged dataset for the two populations.

### Approximate Bayesian computation for inferring hybrid population history

We used a variety of approaches to investigate the time since admixture in the Santa Cruz and Huextetitla hybrid populations, described in detail in Supporting Information 5. However, many approaches assume a single pulse of admixture, which may not be realistic for the Santa Cruz and Huextetitla hybrid populations where hybrids and pure *X. birchmanni* coexist (see Results).

To investigate this, we used an approximate Bayesian computation approach to estimate the history of admixture consistent with observed data in the Santa Cruz and Huextetitla hybrid populations. Guided by results of initial simulations (see Results), we drew parameters from a uniform prior for the time since initial admixture of 10-200 generations, admixture proportion of 0.7-1 *X. cortezi*, and hybrid population size ranging from 50-3,000 diploid individuals. We also implemented migration into the hybrid population from sympatric *X. birchmanni* individuals. Based on the number of early generation hybrids between ancestry clusters observed in the empirical data (see Results), we knew migration rates were low. We thus drew a per-generation migration rate from sympatric *X. birchmanni* individuals of 0-2%. We performed simulations in SLiM (Haller & Messer, 2019) and used the tree sequence recording functions to track individual ancestry (Haller, Galloway, Kelleher, Messer, & Ralph, 2018). All scripts to implement these simulations are available on github (https://github.com/Schumerlab/Xbir_xcor_hybridzone).

To identify the subset of simulations most closely matching patterns in our data, we performed rejection sampling at a 5% threshold based on summary statistics from our data and from simulations. As summary statistics we used average genome-wide ancestry, population-level variance in genome-wide ancestry, and the average length of minor parent ancestry tracts. We performed simulations until 500 parameter sets had been accepted. After an initial set of 1 million simulations resulted in only tens of accepted parameter sets for Huextetitla, we restricted parameter space guided by those accepted to simulate a more restricted range of initial admixture proportions (0.85-1) and migration rates (0-0.5%), and a broader range of generations since initial admixture (10-500). Otherwise simulations for Huextetitla were performed as described above.

### Evaluating evidence for assortative mating in the Santa Cruz hybrid population

Evidence of bimodal ancestry structure in both hybrid populations (see Results) is suggestive of ancestry assortative mating, strong selection on hybrids, or habitat partitioning. To investigate this, we collected 87 females from the Santa Cruz hybrid population in March of 2020, euthanized them, and dissected and developmentally staged their offspring (Supporting Information 6). Forty-six females had developing embryos, with an average of 18 and standard deviation of 10 per female; past work has suggested that a brood typically contains ~3 sires (Paczolt et al., 2015; Schumer et al., 2017). For each brood, we randomly selected two offspring for sequencing from each developmental stage present (to account for possible developmental differences associated with mating type). This resulted in a total of 159 sequenced embryos across mothers (mean 4.4, standard deviation 6.8 per mother), which were used in low-coverage library preparation and sequencing as described above.

To evaluate evidence for assortative mating by ancestry, we took advantage of expectations about maternal-offspring ancestry differences as a function of different types of mating events. Given the extreme differences in ancestry observed across the two genetic clusters in the Santa Cruz hybrid population (Fig. 2), the difference between a mother and her offspring in ancestry allows us to infer the ancestry of the father. Specifically, if a female mates with a male from her own genetic cluster, she and her offspring will have very similar genome-wide ancestry, with the difference between them falling close to zero. If a female instead mates with a male from the other subpopulation, she and her offspring are expected to differ by ~40% in their genome-wide ancestry, given a difference of more than 80% in admixture proportions between the two clusters (Fig. 2). This allowed us to quantify the evidence for assortative mating in observed mating events compared to simulations with varying strengths of assortative mating (Supporting Information 7). We had originally planned to analyze evidence for differential development as a function of mating type, but found too few mating events between ancestry clusters for this analysis to be conducted (Supporting Information 6).

### Analysis of videos from the Santa Cruz hybrid population

As a first step towards evaluating whether there is evidence of habitat partitioning in this structured hybrid population, we took underwater videos from the Santa Cruz hybrid population. Because males of the two clusters can be reliably distinguished based on their morphological characteristics, we scored videos to evaluate whether males were inhabiting the same space.

Underwater video footage was recorded at the Santa Cruz locality to determine whether there is spatial and temporal overlap between *X. birchmanni* and *X. cortezi-*cluster males at this site. Videos were taken consecutively in 50 second to 23 minute sections (20 videos, total of 267 minutes) in July 2020. Cameras were set up in shallow pools isolated by riffles up and down-stream, and the frame of view spanned ~1.5 meters. Males of the two clusters are visually distinguishable by the presence or absence of a sword (see Results), so the number of sworded and unsworded adult males observed was recorded for each video. Each time an adult male swordtail entered the ~1.5 meter frame of view was considered an independent observation and we observed 52 instances of male swordtails entering the frame of view. The presence of sworded and unsworded adult males in the same video was considered evidence for spatial and temporal overlap between the two genetic clusters. Females of the two genetic clusters are not visually distinguishable and thus were not evaluated.

## Results

### Demographic history of X. cortezi and split from X. birchmanni

We used the *X. cortezi* (population Las Conchas) data obtained from 10X sequencing, along with pre-existing sequence data for single individuals of *X. birchmanni* (Coacuilco locality) and *X. cortezi* (nacimiento de Huichihuayán locality, San Luis Potosí; Powell et al., 2020; Schumer et al., 2018), to compare the demographic histories of *X. cortezi* and *X. birchmanni* (see Supporting Information 8). PSMC analysis of each individual indicates distinct demographic histories of *X. birchmanni* and *X. cortezi* populations (Fig. 1; assuming two generations per year and a mutation rate of 3.5 × 10^−9^). Interestingly, our results also suggest divergent demographic trends between two *X. cortezi* populations near the hybrid zone (Las Conchas and nacimiento de Huichihuayán; Fig. S4). Declines in effective population size over the last 20,000 years inferred from the individual sampled from Las Conchas may reflect the demographic effects of colonization of this small tributary (Fig. 1).

Despite differences in the timing of population size fluctuations, the long-term effective population size across species and sampling sites, estimated based on the harmonic mean (Supporting Information 8), was quite similar between the *X. cortezi* population at nacimiento de Huichihuayán and *X. birchmanni*. Specifically, we estimated that the long-term effective population size for *X. cortezi* ranged from 47,000-56,000 across populations compared to 48,000-53,000 in *X. birchmanni,* consistent with the observation that levels of genetic diversity are similar between these *X. cortezi* and *X. birchmanni* populations (0.1% and 0.12% per basepair respectively). Assuming a long-term effective population size of 50,000 for both species, we estimate that *X. cortezi* and *X. birchmanni* diverged from each other approximately 250,000 years ago (Supporting Information 8).

### Santa Cruz and Huextetitla populations are composed of pure X. birchmanni and X. birchmanni x X. cortezi hybrids

Initial analysis of a high-coverage individual sampled from Huextetitla indicated that this individual was a hybrid between *X. birchmanni* and *X. cortezi* (D= −0.49, Z= −30; see also Supporting Information 1), motivating us to develop the local ancestry inference approaches as described in the Methods and Supporting Information 2-4. After inferring local ancestry based on the 1.1 million ancestry informative sites developed for *X. birchmanni* x *X. cortezi* hybrids, we summarized genome-wide ancestry for each individual sampled at the Huextetitla and Santa Cruz locations. To do so, we converted posterior probabilities at each ancestry informative site to hard-calls, requiring that the posterior probability for a given ancestry state exceed 0.9.

This analysis uncovered two genetically distinct subpopulations present in both the Huextetitla and Santa Cruz locations. One cluster consisted of nearly pure *X. birchmanni* with mean *X. cortezi* ancestry of 0.6±0.1% (N=64) and 1±0.6% (N=59) at Huextetitla and Santa Cruz respectively, coexisting with the second cluster of *X. birchmanni* x *X. cortezi* hybrids, with mean *X. cortezi* ancestry of 91±1% (N=12) and 86±6% (N=36) at Huextetitla and Santa Cruz respectively (Fig. 2). These results suggest the presence of strong barriers to gene flow between the two ancestry clusters at both locations, which we explore in more detail below. Notably, given the geography of these river systems (Fig. 1), the two hybrid populations likely formed independently and are presently allopatric.

### Wild X. birchmanni and X. birchmanni x X. cortezi hybrids can be distinguished by their phenotypic differences

Due to morphological differences between species (Fig. 1) and striking differences in ancestry between the two genetic clusters at both the Huextetitla and Santa Cruz sampling sites (Fig. 2), we predicted that males of the two clusters could be distinguished phenotypically. As is the case with many swordtail species, females of the two species are not visually distinguishable (Fig. S5). Using traits that differentiated males in PCA analysis (Fig. 2), we tested how well male genotypes could be predicted based on these phenotypes using discriminant function analysis. We found that a linear discriminant function analysis model fit to 75% of individuals (parental species and hybrids from both sites) accurately predicted the ancestry cluster of 90.9% of individuals not used to fit the model (N = 22). We note that we did not have sufficient individuals to perform training separately on the two sampling sites. Training on different subsets of the data ranging from 30-90% did not qualitatively change results (accuracy range: 89-94%).

### ABC simulations indicate that hybrid populations formed recently

Because the *X. birchmanni* and hybrid *X. cortezi* ancestry clusters are sympatric, we realized that typical approaches to estimate the time of admixture between the two species would likely underestimate the time of initial admixture. As a result, we used an ABC approach allowing for ongoing migration to infer population history. We focused these simulations on the hybrid *cortezi* ancestry cluster, as the *X. birchmanni* cluster shows little evidence of admixture.

From both the Santa Cruz and Huextetitla populations, we inferred well-resolved posterior distributions for the time since initial admixture and migration rates from the *X. birchmanni* cluster into the hybrid *X. cortezi* cluster (Fig. 3). For both Santa Cruz and Huextetitla, we did not recover a well-resolved posterior distribution for population size, but posterior distributions are skewed away from very small population sizes for both sets of simulations (<500 individuals; Fig. S6). Although our simulations allow us to infer initial admixture proportions in the Huextetitla population (Fig. 3), we were surprised that we were unable to recover a well resolved posterior distribution for initial admixture proportion for the Santa Cruz hybrid population. However, this appears to be driven by a strong correlation in the posterior distributions between admixture proportion, time of initial admixture, and migration rate parameters inferred for Santa Cruz. Joint posteriors for these parameters for the Santa Cruz population are shown in Fig. 3 and Fig. S7.

**Figure 3.**
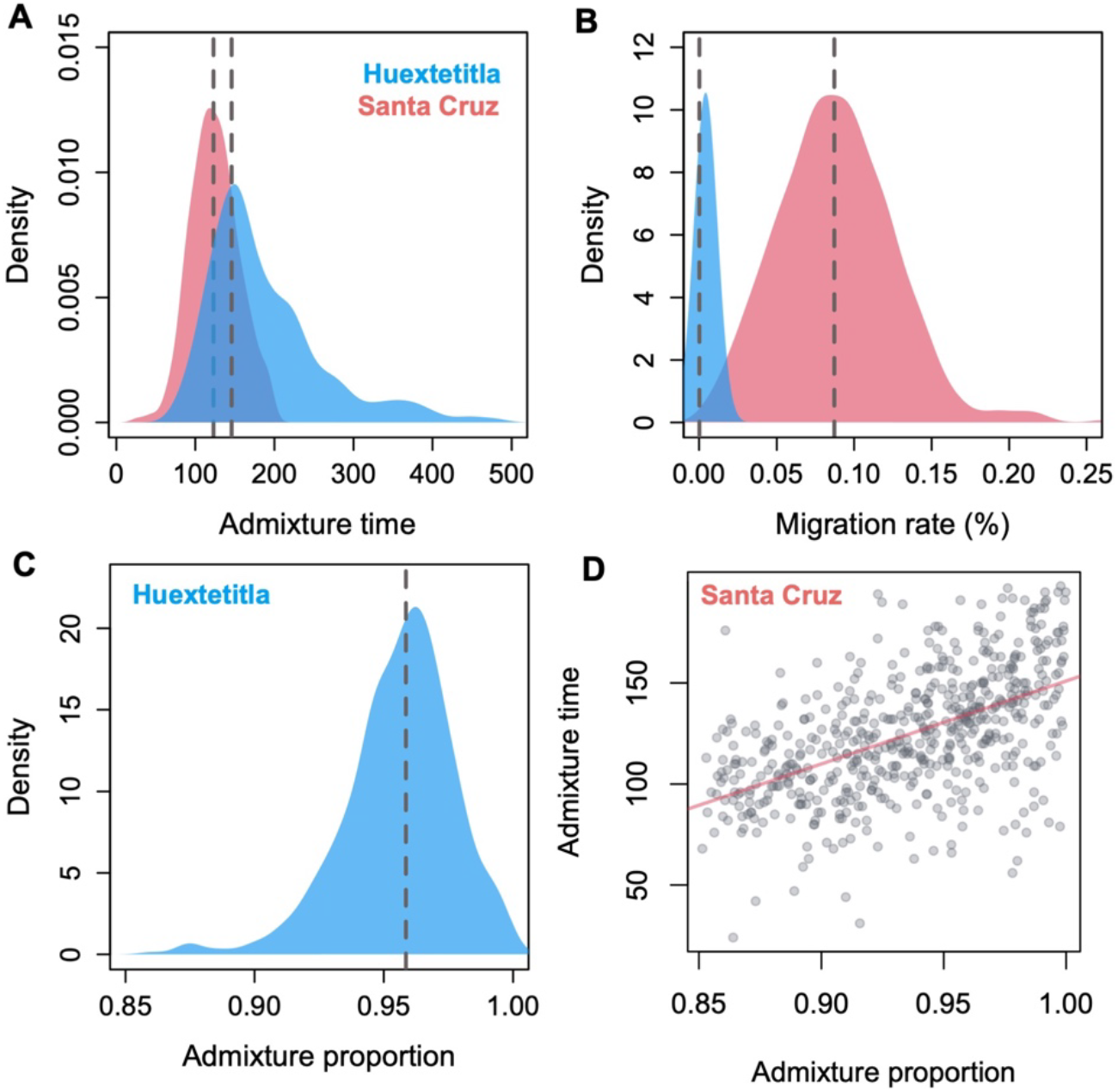
Posterior distributions from Approximate Bayesian Computation simulations inferring demographic history of the *X. cortezi* ancestry cluster in Huextetitla (blue) and Santa Cruz (pink). **A**) Posterior distributions for admixture time indicate that both populations formed relatively recently, in the last ~150 generations. **B**) Posterior distributions of per-generation migration rate reflect substantial differences between populations which can also be observed in variation in admixture proportion (Fig. 2). **C**) For the Huextetitla population, where cross-cluster migration rates are much lower, we recovered a well-resolved posterior distribution of initial admixture proportion. **D**) For the Santa Cruz population, accepted initial admixture proportions span a wide range of parameters and co-vary with both the time since initial admixture (shown here) as well as the cross-cluster migration rate (Fig. S7).

Posterior distributions for admixture time suggest that the hybrid *X. cortezi* populations at Santa Cruz and Huextetitla formed recently, within the last ~140 and ~167 generations respectively (95% confidence intervals – Santa Cruz: 101-183 generations; Huextetitla: 92-384 generations). These estimates are older than estimates from LD decay methods (Supporting Information 5), which put the time of initial admixture ~40 generations ago. This discrepancy is not entirely surprising because LD decay methods tend to underestimate the time since initial admixture in cases where there is ongoing hybridization, and ABC simulations suggest moderate levels of ongoing gene flow from the *X. birchmanni* ancestry cluster into the *cortezi* hybrid ancestry cluster at Santa Cruz (maximum a posteriori or MAP estimate of *m=*0.1%, 95% confidence intervals: 0.02-0.15%). Ongoing migration appears to be much more limited at the Huextetitla collection site (MAP estimate of *m*=0.001%, 95% confidence intervals: 0.0002-0.01%).

Notably, the low inferred migration rates despite the populations existing in sympatry suggests some substantial barrier to gene flow – whether it be genetic, ecological, or via assortative mating. We explore these possible barriers in more detail below.

### Evidence for ongoing admixture and assortative mating

Out of 49 pregnant females collected from the Santa Cruz hybrid population, we successfully sequenced the mother and at least one offspring for 46 mother-offspring pairs. Thirty of these mothers belonged to the hybrid *X. cortezi* genotype cluster and 16 were nearly pure *X. birchmanni*. Based on observed ancestry in embryos, none of the offspring collected were the product of a first generation cross-cluster mating event, however we infer that two females from the *X. cortezi* genotype cluster had mated with males of intermediate ancestry (males with approximately 25% and 55% *X. birchmanni* ancestry respectively). The proportion of sampled individuals with intermediate ancestry did not differ between the embryonic and adult populations (4.3±3% of sampled embryos and 3.2±2% of sampled adults with ancestry between 5-75% *X. cortezi*). Notably, all these individuals are inferred to have a *X. cortezi* mother, which could hint at weaker assortative mating by ancestry among females of the *cortezi* cluster (Supporting Information 9). We found no evidence of differences in number of embryos, variation in embryo stage, or developmental abnormalities between females of the two clusters (all p>0.7; Supporting Information 6).

Analysis of the maternal-offspring ancestry patterns indicate clear deviations from expectations under random mating (Fig. 4; Supporting Information 7). We used simulations to quantify the strength of ancestry-assortative mating consistent with our data (Fig. 4B, Supporting Information 7). These simulations indicated that our data is consistent with a strength of ancestry assortative mating of approximately 98% (Fig. 4C). Strong assortative mating by ancestry is thus one likely factor maintaining the two distinct subpopulations at Santa Cruz. We note that our current results do not allow us to distinguish between assortative mating mediated via mate preferences and other possibilities such as near-perfect sperm precedence for males of similar ancestry or nearly complete mortality of cross-cluster offspring in the earliest stages of embryonic development.

**Figure 4.**
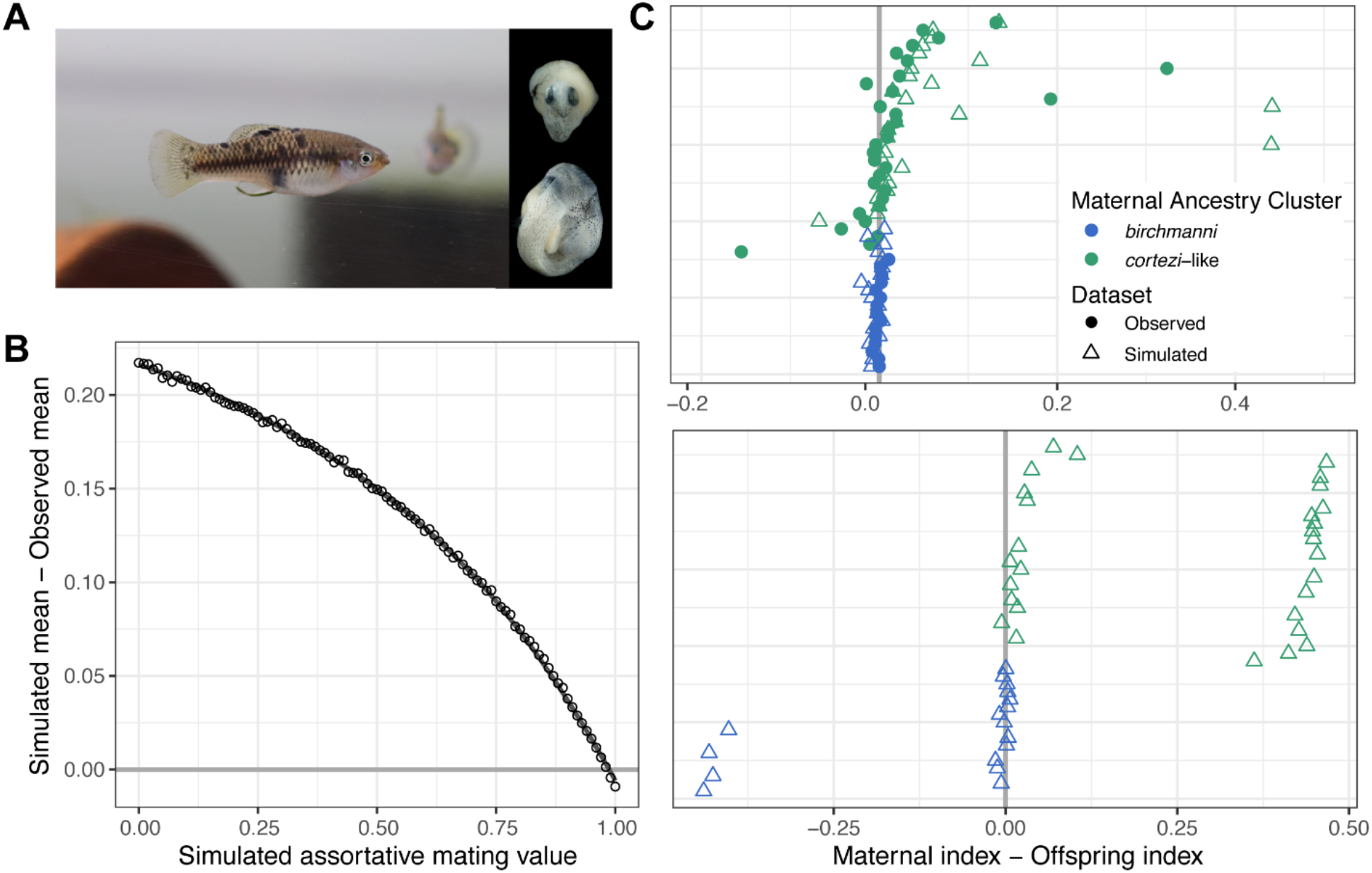
Results of assortative mating simulations in the Santa Cruz population. **A)** Photos of a pregnant *X. cortezi*-cluster female and embryos. **B)** Results of simulations ranging from 0-100% assortative mating in increments of 1% comparing the simulated versus observed difference in maternal and offspring ancestry index. Simulations of 98% assortative mating minimized the difference between the observed and simulated datasets. **C)** In observed (circles) and 98% assortative mating simulated (triangles) datasets showing the difference between the maternal and offspring ancestry index (top), few offspring have dramatically different ancestry from their mothers. In contrast, many such individuals are observed in simulations of random mating (bottom). Points close to the zero line represent females that mated with males from their own ancestry cluster. Individuals are colored based on their maternal ancestry cluster and are placed on the y-axis based on increasing *X. cortezi* ancestry.

### Evidence of sympatry of the X. birchmanni and cortezi populations

Ancestry assortative mating could be driven by processes such as mate discrimination or by spatial isolation that prevents individuals from different genotype clusters from encountering each other. We suspected that the latter scenario was not the case at these collection sites as we repeatedly collected both male and female *X. birchmanni* and *X*. *cortezi* cluster hybrid individuals from the same minnow traps (over three collections at Santa Cruz and one collection at Huextetitla). This suggests that these individuals are sympatric in the wild and have the opportunity to mate with each other.

As a first step towards investigating this further, we took underwater videos during the summer of 2020 at Santa Cruz and scored the videos for interactions between males of the two ancestry clusters, which can be distinguished with high accuracy based on their sword phenotypes (see above). We found both sworded and unsworded males in 4 of the 12 videos in which male swordtails were observed (12 to 15 minutes each, total of 158 minutes), showing that individuals of both hybrid clusters inhabit the same areas at the same time. There were 8 videos (50 seconds to 23 minutes each, total of 109 minutes) in which no male swordtails were observed. Raw video footage and scored data are available on Dryad (Accession: XXXXX).

## Discussion

In the past two decades we have found that hybridization occurs much more often than previously thought, and have made phenomenal progress characterizing the frequency of hybridization between species across the tree of life. One of the next frontiers in hybridization research is understanding the extent to which the evolutionary outcomes of hybridization are predictable across pairs of species, from the genetic to the population level.

Here, we develop sensitive local ancestry calling and infer the history of hybridization and ancestry structure in two newly characterized hybrid populations between non-sister *Xiphophorus* species (Fig. 2). *X. birchmanni* and *X. cortezi* are more distantly related than sister species *X. birchmanni* and *X. malinche*, which have become an emerging model system for studying the consequences of hybridization between species (Fig. 1). Notably, like the *X. birchmanni* x *X. malinche* system, our demographic inference suggests that these hybrid populations formed recently (in the last ~150 generations; Fig. 3), providing a window into evolution in the earliest stages after hybridization.

Given that the Santa Cruz and Huextetitla populations appear to be geographically independent—as they occur in two separate rivers—the similarities in overall ancestry structure and inferred demographic parameters between the populations are striking. While the ancestry structure of the populations appears to be driven by strong assortative mating (see below), the concordance in demographic parameters is more puzzling. This could indicate that the Santa Cruz and Huextetitla sites are not as isolated as their current geography would suggest (Fig. 1). Alternately, the concordance in certain parameters, such as the time since initial admixture, could reflect shared histories of disturbance due to their geographical proximity to growing human settlements, as appears to be the case in the *X. birchmanni* x *X. malinche* hybrid zones (Fisher et al., 2006). Dense sampling along clines in these two rivers will help us distinguish these possibilities.

The existence of distinct ancestry clusters in both populations suggests substantial reproductive barriers. While the simplest explanation for this pattern would be some form of spatial isolation by ancestry, several observations argue against this explanation. First, we collected reproductively active males and females of both ancestry clusters in the same minnow traps over multiple collections. Second, we identify males of both clusters in underwater videos capturing small geographic areas. Instead, the evidence argues for a strong role of assortative mating in driving ancestry structure. Based on sequencing of wild-caught mothers and their offspring, we find strong evidence for nearly complete assortative mating by ancestry cluster.

Despite strong ancestry assortative mating, the results of ABC simulations are consistent with low levels of ongoing gene flow between ancestry clusters (Fig. 3). Intriguingly, our data show that all individuals originating from cross-cluster mating events had mothers from the *cortezi* ancestry cluster. This hints that mating barriers may be weaker between *X. cortezi* females and *X. birchmanni* males than in the alternative direction, consistent with higher levels of *X. birchmanni* ancestry in the *cortezi* ancestry cluster (Fig. 2). Alternately, these results could be explained by asymmetric genetic barriers such as embryonic lethality in the earliest stages of development, since we only sequenced embryos that were visually identifiable as fertilized (i.e. post-blastodisc phase). Indeed, selection against hybrid ancestry is probable regardless of the strength of assortative mating in the two clusters, given that the parental species are more distantly related than *X. birchmanni* and *X. malinche*, which have well-documented genetic incompatibilities (Powell et al., 2020; Schumer et al., 2014). Teasing apart the relative contributions of different barriers to gene flow in *X. birchmanni* × *X. cortezi* hybrid populations will be an exciting avenue for future work.

Perhaps the most striking finding of this study is the repeatability of ancestry structure across diverse hybrid zones. The bimodal population structure we observe is repeated not only between the Santa Cruz and Huextetitla populations of *X. birchmanni* x *X. cortezi* hybrids but has also been found in previously studied *X. birchmanni* × *X. malinche* hybrid populations. The ancestry structure of the *X. birchmanni* x *X. cortezi* populations mirrors that of the *X. birchmanni* x *X. malinche* hybrid population on the Río Calnali (“Aguazarca”). This population consists of a cluster of *birchmanni*-skewed hybrids deriving ~75% of their genome from *X. birchmanni* and introgressed *X. malinche* individuals, deriving ~5% of their genome from *X. malinche* (Culumber, Ochoa, & Rosenthal, 2014; Schumer et al., 2017). Moreover, strong ancestry assortative mating also maintains isolation between ancestry clusters in this hybrid population (Culumber et al., 2014; Schumer et al., 2017).

The repeatability of these patterns across distinct hybridizing species pairs highlights the importance of assortative mating in shaping ancestry and population structure in these young hybrid populations. We might expect that at least some of the mechanisms driving assortative mating are shared across systems, since *X. birchmanni* females are known to prefer swordless males (Wong & Rosenthal, 2006). However, behavioral mating preferences have been difficult to detect in *X. birchmanni* x *X. malinche* hybrids. Investigating the extent to which factors that generate or disrupt assortative mating are shared across the *Xiphophorus* phylogeny will be another rich area for future study.

The web of hybridization between *X. cortezi, X. birchmanni*, and *X. malinche* provides a novel opportunity to investigate the consequences of hybridization across scales. Greater sample sizes across both *X. birchmanni* x *X. cortezi* populations will allow us to test shared drivers of local ancestry across systems (Schumer et al., 2018), identify hybrid incompatibilities, and ask whether observed patterns are a function of phylogenetic history or other biological variables. Such comparative approaches, made possible by the work described here, will ultimately allow us to evaluate the degree to which outcomes of hybridization are predictable across independent hybridization events.

## Acknowledgements

We thank Gil Rosenthal, Andrea Sweigart, Vaclav Alexei Sotola, Matthew Farnitano, Kira Delmore, Yaniv Brandvain, and members of the Schumer lab for helpful discussion and/or feedback on earlier versions of this work. We also thank Baruc Zago-Mazzocco for field work support. We are grateful to the Mexican federal government for permission to collect samples. We thank Stanford University and the Stanford Research Computing Center for providing computational support for this project. This work was supported by NSF GRFP 2019273798 to B. Moran, NRSA F32 GM135998 to B. Kim, a Cornell University Provost Diversity Fellowship to S. M. Aguillon, a CEHG fellowship and NSF PRFB (2010950) to Q. Langdon, and a Hanna H. Gray fellowship, NIH 1R35GM133774, and Human Frontiers in Science (RGY0081) grant to M. Schumer.

