## Supporting Information for "Two new hybrid zones expand the swordtail hybridization model system"

**Supporting Information 1. SppIDer analysis of ancestry mixture proportions**

As an initial exploration of potential contributions from *X. birchmanni*, *X. cortezi*, and *X. variatus*, which have all been found near the hybrid populations at Huextetitla and Santa Cruz, we used sppIDer (Langdon, Peris, Kyle, & Hittinger, 2018), a competitive mapping and depth analysis pipeline. Reads from a deep-sequenced Huextetitla individual were mapped to a combination reference genome that consisted of the *X. birchmanni* reference genome and the *X. cortezi* and *X. variatus* reference genomes from the Ragout output of the cactus alignment described in the main text. All genomes were hard masked for repetitive elements.

We found that 75.4% of the reads preferentially mapped to the *X. cortezi* reference genome, 22.32% mapped to the *X. birchmanni* reference genome, while only 2.28% of the reads mapped to the *X. variatus* reference genome (Fig. S8). This analysis suggested that this individual is a hybrid between *X. birchmanni* and *X. cortezi*, derives the majority of its genome from *X. cortezi*, and has little contribution from *X. variatus*.

**Supporting Information 2. Local ancestry inference based on three source populations**

Although SppIDer results on deep-sequenced individuals were not suggestive of hybridization between *X. variatus* and northern swordtails in the Huextetitla population, out of an abundance of caution, we also evaluated this in our population samples. To do so, we took advantage of AncestryHMM's ability to model three source populations in local ancestry inference (Corbett-Detig & Nielsen, 2017).

Building off of ancestry informative sites identified between *X. birchmanni* and *X. cortezi*, we added sites that differentiated *X. variatus* from *X. birchmanni* and *X. cortezi*. In contrast to datasets available from *X. birchmanni* and *X. cortezi*, we did not have large numbers of available parental individuals that had been re-sequenced (five individuals from two populations). However, due to deep divergence between the northern swordtail clade and *X. variatus* we expect low levels of incomplete lineage sorting and a high density of ancestry informative sites. Indeed, after filtering sites that were not fixed in the five *X. variatus* individuals our dataset included 4.5 million ancestry informative sites that differentiated *X. variatus* from the two northern swordtail species (~6 per kb).

Result of three-way local ancestry calling were consistent with those including only two source populations. Specifically, few individuals had evidence for any *X. variatus* ancestry, with genome-wide estimates ranging from 0-2% (median *X. variatus* ancestry of 0). The only exception was an *X. variatus* juvenile from Huextetitla, who was falsely assigned phenotypically as *X. birchmanni*, and was inferred to have 100% *X. variatus* ancestry (Fig. S9).

**Supporting Information 3. Simulations of drifted reference populations**

AncestryHMM relies on allele frequency counts at ancestry informative sites that distinguish the parental species (Corbett-Detig & Nielsen, 2017). However, accuracy is expected to be lower when reference panels from the appropriate source populations are not available (Schumer, Powell, & Corbett-Detig, 2020), because drift between the hybridizing and reference populations that alter allele frequencies at ancestry informative sites can contribute to errors in local ancestry inference. We were particularly concerned about this since both the *X. birchmanni* and *X. cortezi* allopatric source populations are quite geographically isolated from the hybrid populations (Fig. S4).

To explore this in simulations, and evaluate what steps we could take to reduce such errors, we used a simulator we recently developed called *mixnmatch* (Schumer et al., 2020) to simulate genetic drift between the source and hybridizing *X. birchmanni* and *X. cortezi* populations. To define ancestry informative sites in simulations we sampled from populations that diverged from the true source populations 0.25*Ne* generations ago. We then used *mixnmatch* to simulate admixture 150 generations before the present with mixture proportions matching those observed in the Santa Cruz population. We used the local recombination map from *X. birchmanni* for chromosome 1 in simulations and simulated a 10 Mb chromosome for 50 individuals.

We found that under a scenario with drift between the reference and source populations, per-site error rates were approximately 1%, compared to 0.3% in simulations without drift. This indicates that, as expected, the HMM is sensitive to the allele frequency estimates in the source populations. Because we do not have access to additional parental populations for our real data, we next explored how we might mitigate this issue.

Examining errors in the simulated data, we found that they were most common at sites inferred to be completely fixed in the drifted reference populations that were segregating in the true source population. We reasoned that by defining the minor allele frequency as small instead of fixed, we might improve accuracy. For each fixed ancestry informative site, we added one allele count to the alternate allele in each species, re-ran the HMM, and summarized accuracy. We found that this simple modification reduced our error rate by three-fold. We thus proceeded with this approach for the real data.

**Supporting Information 4. Simulations to test the sensitivity of priors in the HMM**

AncestryHMM requires input parameters including generations since initial admixture, admixture proportions, and error rate. Because some of these parameters for *X. birchmanni* x *X. cortezi* hybrid populations are unknown, or could be estimated with error, we wanted to perform simulations to evaluate how sensitive the HMM is to user-provided priors for these values.

To explore this, we performed simulations with the programs *mixnmatch* and *ancestryifer* where the true admixture time was much older (5X) or younger (1/5) that the user-provided prior admixture time (150 generations since initial admixture). We repeated this same procedure for error rates, using *mixnmatch* to simulate contamination as a way to introduce error. In these simulations, 2.5% of the reads per individual were drawn from other simulated individuals, and the HMM was run with prior error rates of 0%, 2.5% and 5% specified. We did not perform simulations evaluating admixture proportion priors since we are able to accurately estimate this value from the raw data (see main text).

We found that in simulations where the prior admixture time was set as much younger (30 generations since initial admixture) than the true value, error rates were comparable to simulations with the correct prior (error rate of 0.2% per ancestry informative site). When the prior admixture time was set to 750 generations (5X the true value), error rates did increase, but modestly (0.4%). This suggests that although error rates can increase with misspecification of the admixture time, presumably through an increase in switch errors, the HMM implemented through AncestryHMM is not very sensitive to admixture time misspecification.

Misspecifying the error rate as 0% in the presence of contamination also led to a slight increase in the error in ancestry assignment (0.4% per ancestry informative site). However, overestimating the error rate or providing the correct error rate resulted in similar accuracy as above (0.1-0.2%). Again, these results suggest that AncestryHMM performs relatively well in the presence of misspecified parameters but highlights some sensitivity to overestimating generations since admixture and underestimating the error rate.

**Supporting Information 5. Inference of admixture time from AncestryHMM and LD decay-based methods**

The timing of admixture is a key parameter in understanding the population history of *X. birchmanni* x *X. cortezi* hybrid zones. In the main text we describe an ABC based approach using forward-time simulations of hybrid populations and ancestry tract lengths to estimate an initial admixture time of ~150-200 generations before the present (Fig. 3).

We complemented this approach with two other methods for estimating admixture time. If users do not specify an admixture time prior, AncestryHMM jointly infers the most likely demographic history consistent with the local ancestry data including time of admixture. Although this approach is sensitive to population demography, it has been reported to be relatively robust to different demographic scenarios at least in the case of recent admixture (Corbett-Detig & Nielsen, 2017). Applying this approach, AncestryHMM inferred a time of initial admixture of 69 generations before the present for the Santa Cruz population.

We also used the decay in ancestry linkage disequilibrium (LD) over genetic distance as a second approach to estimate the time of admixture. This method assumes a single pulse of admixture, which may not appropriately model the history of admixture in either population, since we find evidence for low levels of ongoing admixture with sympatric *X. birchmanni* (see main text). In cases of multiple pulses of admixture or continuous migration, LD decay methods tend to underestimate the time since initial admixture.

To estimate admixture LD decay, we first thinned our ancestry informative markers to retain one marker every 50 kb. We converted this thinned data to plink format and calculated pairwise LD between all markers within 10 Mb on the same chromosome. Physical distance between markers was converted to genetic distance using a previously developed recombination map for *X. birchmanni* (Schumer et al., 2018). Following previously described methods, we binned data into 0.1 cM windows and fit an exponential curve to the decay in admixture LD over genetic distance. We used the relationship:

$$E(D) = ae^{-Tx} + c$$

where D is ancestry LD, T is the number of generations since initial admixture, x is genetic distance in Morgans, *c* is a constant describing the value to which D decays, and *a* is a coefficient. Values for *a,* T, and *c* were predicted using the non-linear least squares function in R (`nls`) using observed D and inferred genetic distance. This yielded an estimate of 34 ± 2 generations since initial admixture in the Santa Cruz population and 36 ± 3 in Huextetitla. We interpret these more recent estimates of admixture time as a result of the fact that our focal populations violate the assumption of a single pulse of admixture between species.

**Supporting Information 6. Description of embryo staging**

In order to explore any possible relationship between ancestry and developmental abnormality, embryos from the Santa Cruz hybrid population were categorized by developmental stage prior to genotyping. Ovaries were removed from the 79 ethanol-preserved gravid Santa

Cruz females and processed in order of increasing distance from the urogenital opening. Each egg was separated from the ovarian tissue and assigned to a developmental category from 1-11 based on the criteria of (Haynes, 1995); unyolked eggs (stage 1) were not counted or included in further analysis. Embryos which showed characteristics of two developmental categories were assigned the average of the two stages, and those that showed transgressive characteristics (e.g. embryos with markedly little three-dimensional structure given their length) were noted as abnormal and assigned the stage most consistent with their characteristics. Representative photographs of each developmental stage and of any abnormal individuals submitted for sequencing were taken using an AmScope MU1803 18MP camera mounted to an AmScope 3.5X-90X trinocular stereomicroscope.

We summarized the brood characteristics of each mother as the number of eggs carried, the number of stages present, the variance in developmental stage between embryos, and the number of embryos noted as abnormal. We tested *X. birchmanni* and *X. cortezi* cluster mothers for systematic differences in each metric using Welch's *t*-test, and repeated the analysis considering only fertilized eggs (stage > 3). All tests showed no significant difference between the clusters ($P > 0.05$).

**Supporting Information 7. Simulations to evaluate assortative mating by ancestry**

We collected 79 females from the Santa Cruz hybrid population of which 46 had fertilized eggs. To evaluate the level of ancestry assortative mating that best described our data, we performed simulations in R using the paired ancestry of mothers and one randomly chosen offspring. Specifically, we varied the strength of assortative mating and asked how the ancestry differences between mothers and their offspring differed from the observed dataset.

For each simulation, we used the observed maternal and population-level (males and females) ancestries from Santa Cruz as input. We define the "ancestry threshold" as the maximum difference in ancestry between individuals of the same genotype cluster (Fig. 2). We estimated this value from a random normal distribution generated using the observed mean and standard deviation of ancestry of the *cortezi*-like cluster. The probability that a female will accept a mate with a difference in genome-wide ancestry greater than the ancestry threshold is defined as $p_{assortative}$. We used the following procedure in each simulation:

1) For each female, we drew a random ancestry value for her potential mate from the population-level ancestry distribution.
    a. If the difference in ancestry between the female and the simulated mate was less than the ancestry threshold, the simulated mate was accepted.
    b. If the difference in hybrid index was greater than the ancestry threshold, we drew from a random binomial with probability equal to $1\text{-}p_{assortative}$ to determine if the female accepted the simulated mate.
    c. This process was repeated until a mate was accepted.
2) For each pair of mated individuals, we calculated the expected ancestry of their offspring. To add variance in the offspring ancestry to account for the fact that individuals do not inherit exactly 25% of their genome from each grandparent, we drew from a normal distribution with variance equal to the observed variance in ancestry between siblings in the full dataset.
3) We repeated this procedure 500 times for each value of $p_{assortative}$.

We iterated through increasing strengths of $p_{assortative}$ starting at 0 (completely random mating) and increasing by 0.01 in each subsequent simulation up to 1.0 (complete ancestry assortative mating). For each simulation, we then compared the mean and 95% confidence interval of the ancestry differences between simulated mothers and offspring to that of the observed dataset. Based on these simulations, we inferred that the strength of assortative mating that best mimicked patterns in the empirical data from the Santa Cruz hybrid population was 98% ancestry-assortative mating.

**Supporting Information 8. Demographic history of *X. cortezi***

PSMC analysis (Li & Durbin, 2011) to infer the demographic history of *X. cortezi* was carried out using several *X. cortezi* individuals. We used the 10X Chromium library generated for the *X. cortezi* genome assembly (Las Conchas population) as well as previously sequenced *X. cortezi* individuals from Nacimiento de Huichihuayán and *X. birchmanni* individuals from Coacuilco (see main text). Raw reads were processed to remove barcodes and mapped to the *X. birchmanni* reference assembly (Powell et al., 2020) using bwa (Li & Durbin, 2009). Mapped reads were de-duplicated with Picard Tools, and indels were realigned with GATK v3.4 (McKenna et al., 2010). Variants were called using GATK's HaplotypeCaller and GenotypeGVCFs in GVCF mode (McKenna et al., 2010). Of these variants, those with genotype quality GQ > 20, mapping quality MQ > 40, quality-by-depth QD > 10, Fisher strand bias FS < 10, strand odds ratio SOR < 4, mapping quality rank sum MQRankSum > -12.5, read position rank sum ReadPosRankSum > -8, and read depth between 0.5-2X of the genome wide average coverage were retained. Variant sites within 5 bp of an indel were excluded. Likewise, invariant sites were filtered for reference genotype quality score RGQ > 20, and read depth between 0.5-2X of the genome wide average coverage. These hard calls were input to seqtk's mutfa function (https://github.com/lh3/seqtk) to edit the *X. birchmanni* reference, generating a *X. cortezi* pseudo-reference with coordinates identical to *X. birchmanni* genome. Sites that failed the quality metrics described above were masked in the pseudo-reference sequences.

These pseudo-reference FASTA files were converted to a FASTQ with uniform Q-scores of 40, then converted to PSMC input format using fq2psmcfa (https://github.com/lh3/psmc) with default parameters. PSMC analysis was performed on the 24 syntenic *Xiphophorus* chromosomes with the time segmentation parameter set to 4+25*2+4+6 and otherwise default parameters. Estimates of the time to most recent common ancestor (TMRCA) from PSMC were rescaled to $N_e$ using a mutation rate of $3.5\text{x}10^{-9}$ $\text{bp}^{-1}$ $\text{generation}^{-1}$ and a generation time of 0.5 years, as in Schumer et al. (2018). In the case of the individual sampled from the Las Conchas population, where only one sample was available, uncertainty in PSMC estimates was visualized with 100 bootstrap replicates, each constructed by resampling with replacement from non-overlapping 500 kb segments of the genome. Based on variation between bootstraps in estimates of *Ne*, we limited analysis of PSMC results to the past 2,000-100,000 years (Fig. 1). Within this interval, long-term $N_e$ was estimated by the harmonic mean of the $N_e$ estimated by PSMC at each time block, weighted by its length in years.

This data was also used to estimate the divergence time between *X. cortezi* and *X. birchmanni* under a strictly allopatric model. The split time was estimated from the relationship $T_{div(4N)=}0.5(D_{xy}/\theta - 1)$, where θ is the population mutation rate and $D_{xy}$ is the average pairwise divergence between individuals (here the Las Conchas *X. cortezi* and the *X. birchmanni* reference individual). We assumed θ = 0.001 per site for all species, which corresponds to the estimated θ in most swordtail species studied to date including most *X. birchmanni* and *X. cortezi* populations (Schumer et al., 2013; Schumer, Cui, Powell, Rosenthal, & Andolfatto, 2016), and thus is a reasonable estimate of the ancestral θ for these species. We note that this estimate likely represents an underestimate of the divergence time, since $D_{xy}$ between the two species should be decreased by the inferred historic admixture between *X. birchmanni* and *X. cortezi* (Cui et al., 2013).

**Supporting Information 9. Inference of maternal subpopulation ancestry in cross-cluster individuals**

We identified no individuals with intermediate ancestry in our sampling of the Huextetitla population. However, in both our population sampling from the Santa Cruz population and our sequencing of mother-embryo pairs, with identified several individuals with intermediate ancestry. We arbitrarily define intermediate ancestry individuals as those with a proportion of the genome derived from *X. cortezi* falling between 5% and 75%. Four individuals from our population sample of adults fell within this ancestry range, as did three embryos derived from two independent mating events.

Notably, within ancestry clusters, mitochondrial ancestry is fixed for the major parent species (Fig. S10). To determine the maternal cluster of origin for individuals with intermediate ancestry, we extracted ancestry calls at mitochondrial markers. We found that all seven intermediate ancestry individuals had maternal mitochondrial haplotypes that matched the *X. cortezi* ancestry cluster. This observation is unexpected by chance based on the observed admixture proportions of these individuals ($p<0.001$ by simulation). This finding is also consistent with the observation that the *X. birchmanni* subpopulation has extremely low *X. cortezi*-derived ancestry, whereas the *cortezi* subpopulation has substantial ancestry derived from *X. birchmanni*.

We note that these findings could suggest the presence of stronger assortative mating by ancestry among females within the *X. birchmanni* subpopulation, or strong asymmetric genetic barriers. However, such genetic barriers would need to act at the earliest stages of embryonic development to be consistent with our results.

### Supporting Information Tables

**Table S1.** Principal Component Analysis results for phenotypes of males from the Huextetitla and Santa Cruz populations compared to male phenotypes of parental species.

| Population | Principal Component | Percent of variation explained | body length loading | body depth loading | peduncle depth loading | dorsal fin length loading | dorsal fin height loading | sword length loading |
|---|---|---|---|---|---|---|---|---|
| **HUEX** | PC 1 | 0.779 | 0.80 | 0.34 | 0.77 | 0.28 | 0.29 | -0.30 |
| | PC 2 | 0.119 | 0.23 | 0.14 | -0.43 | 0.13 | -0.32 | 0.49 |
| | PC 3 | 0.076 | 0.13 | -0.92 | -0.47 | -0.95 | 0.35 | 0.82 |
| | PC 4 | 0.014 | 0.23 | - | - | - | -0.83 | - |
| | PC 5 | 0.009 | 0.31 | - | - | - | - | - |
| | PC 6 | 0.002 | 0.39 | - | - | - | - | - |
| **STAC** | PC 1 | 0.753 | 0.85 | 0.35 | 0.15 | 0.28 | 0.20 | 0.82 |
| | PC 2 | 0.164 | 0.32 | -0.93 | 0.52 | -0.96 | -0.22 | 0.55 |
| | PC 3 | 0.058 | 0.23 | - | -0.68 | - | 0.48 | - |
| | PC 4 | 0.017 | 0.33 | - | -0.49 | - | -0.81 | - |
| | PC 5 | 0.006 | - | - | - | - | 0.11 | - |
| | PC 6 | 0.003 | - | - | - | - | 0.12 | - |

### Supporting Information Figures

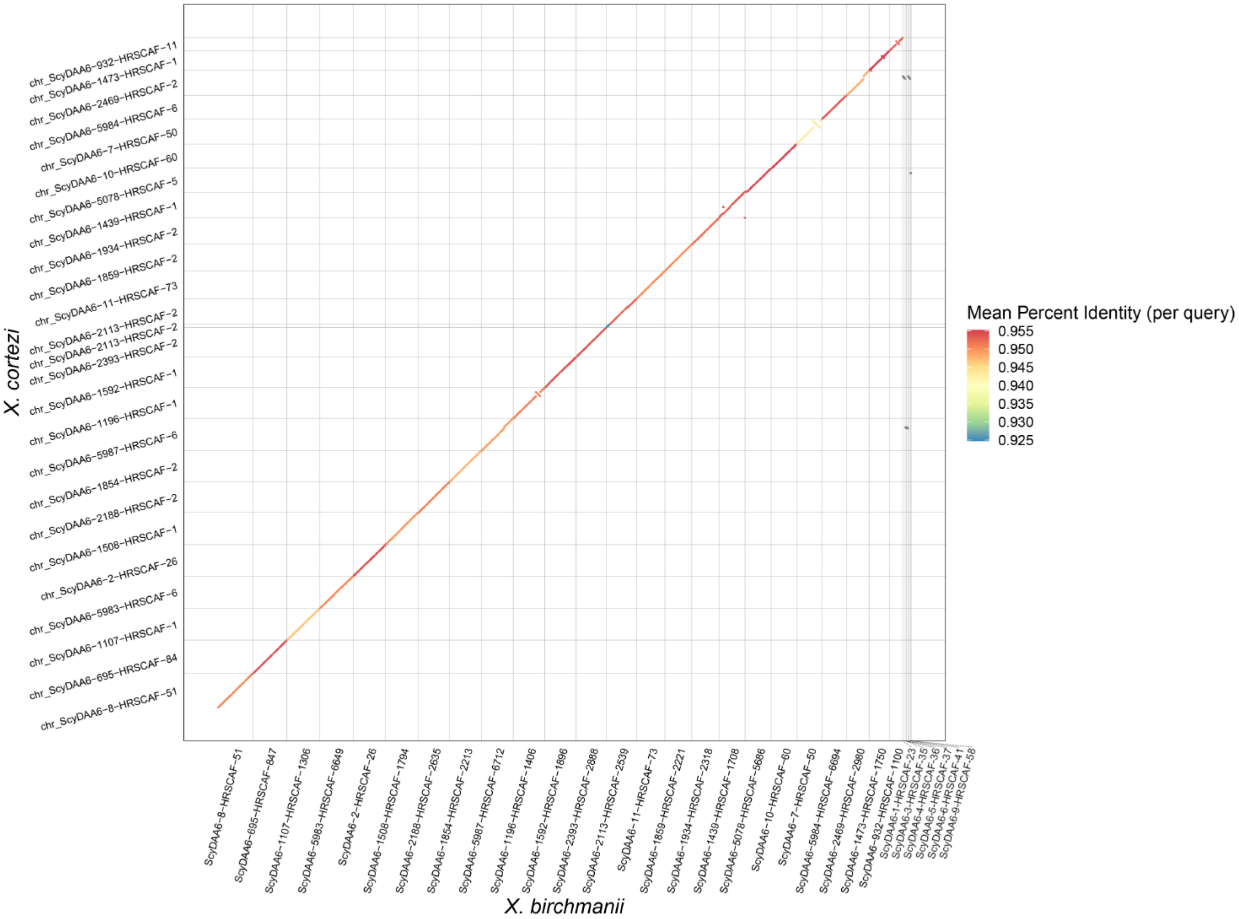


**Figure S1.** Alignment of reference guided *X. cortezi* assembly to the *X. birchmanni* genome. Alignments of less than 50 kbp and *X. cortezi* sequences with total alignments less than 500 kb are not visualized.

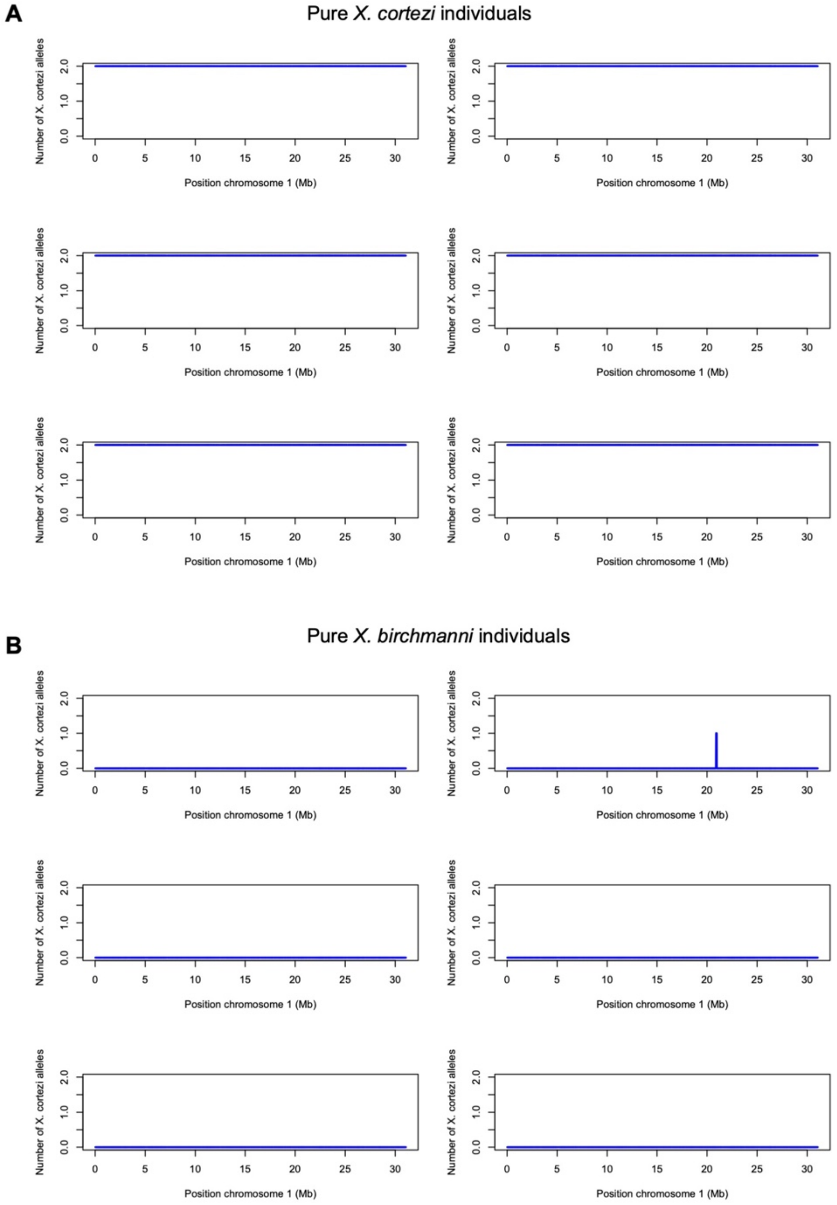


**Figure S2.** Local ancestry on chromosome 1 in a representative sample of pure *X. birchmanni* and *X. cortezi* individuals that were not used to define ancestry informative sites.

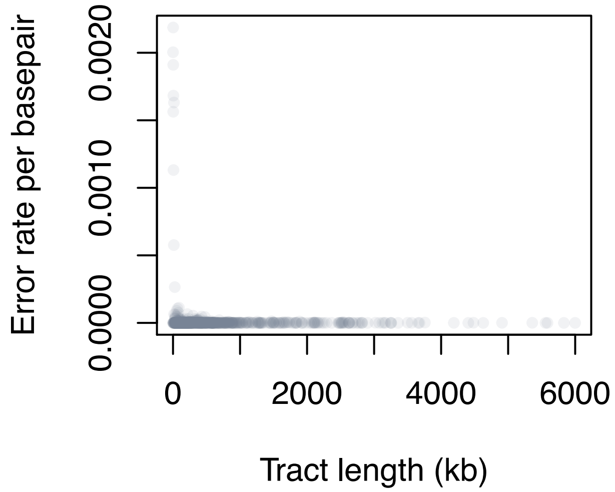


**Figure S3. Accuracy of local ancestry inference based on simulations.** Shown here is the output of simulations with *mixnmatch* showing the per ancestry informative site error rate for each ancestry tract simulated across 50 hybrid individuals. Error rates are below ~0.2% even for short ancestry tracts where error rates tend to be higher. These results suggest that we expect to have high accuracy in local ancestry inference in the Huextetitla and Santa Cruz hybrid populations.

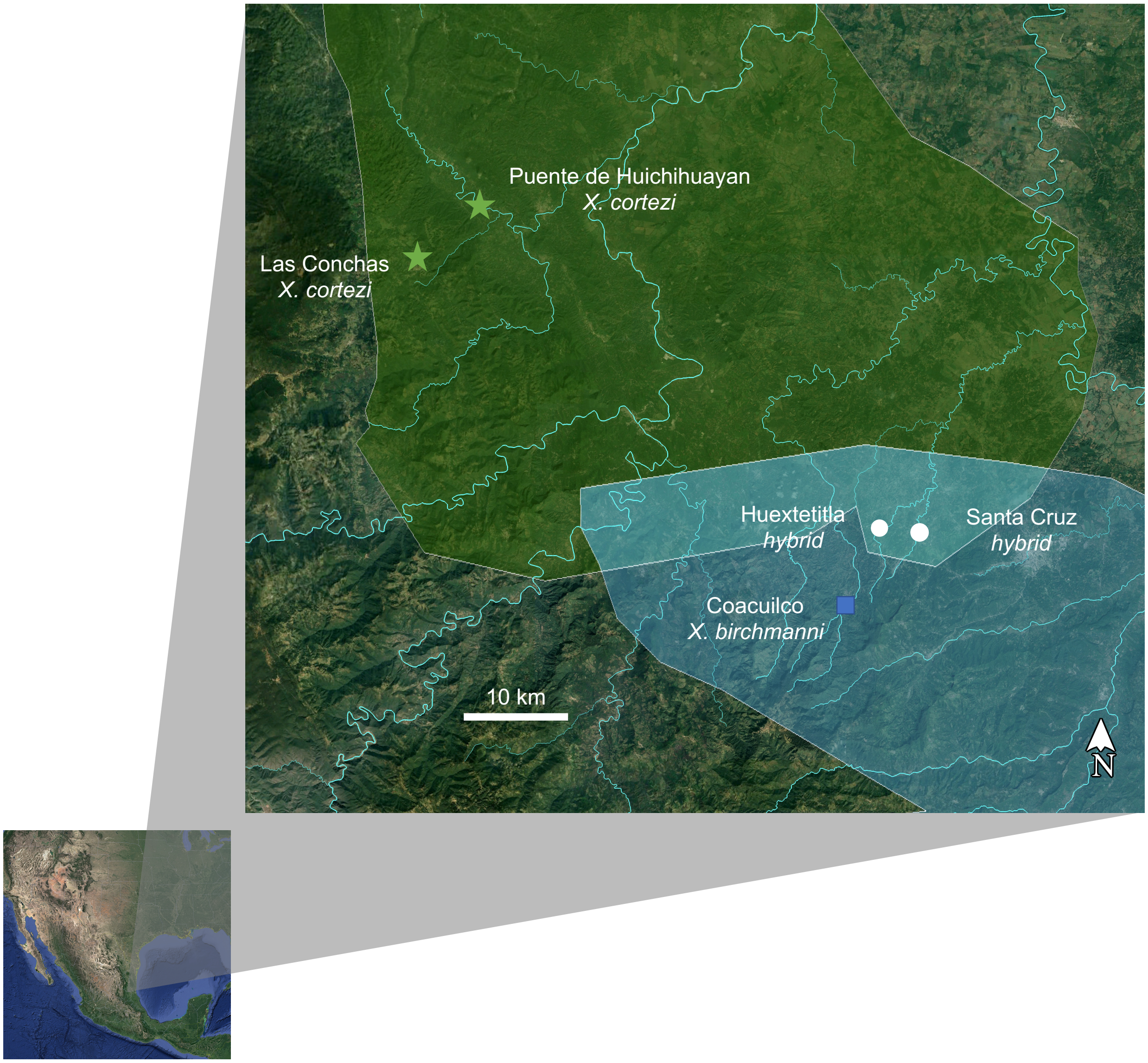


**Figure S4. Map of parental ranges and population collections.** Polygons represent the known ranges of *X. cortezi* (green) and *X. birchmanni* (blue). Pure populations of *X. cortezi* (green stars), *X. birchmanni* (blue square) and hybrid populations (white circles) are labeled.

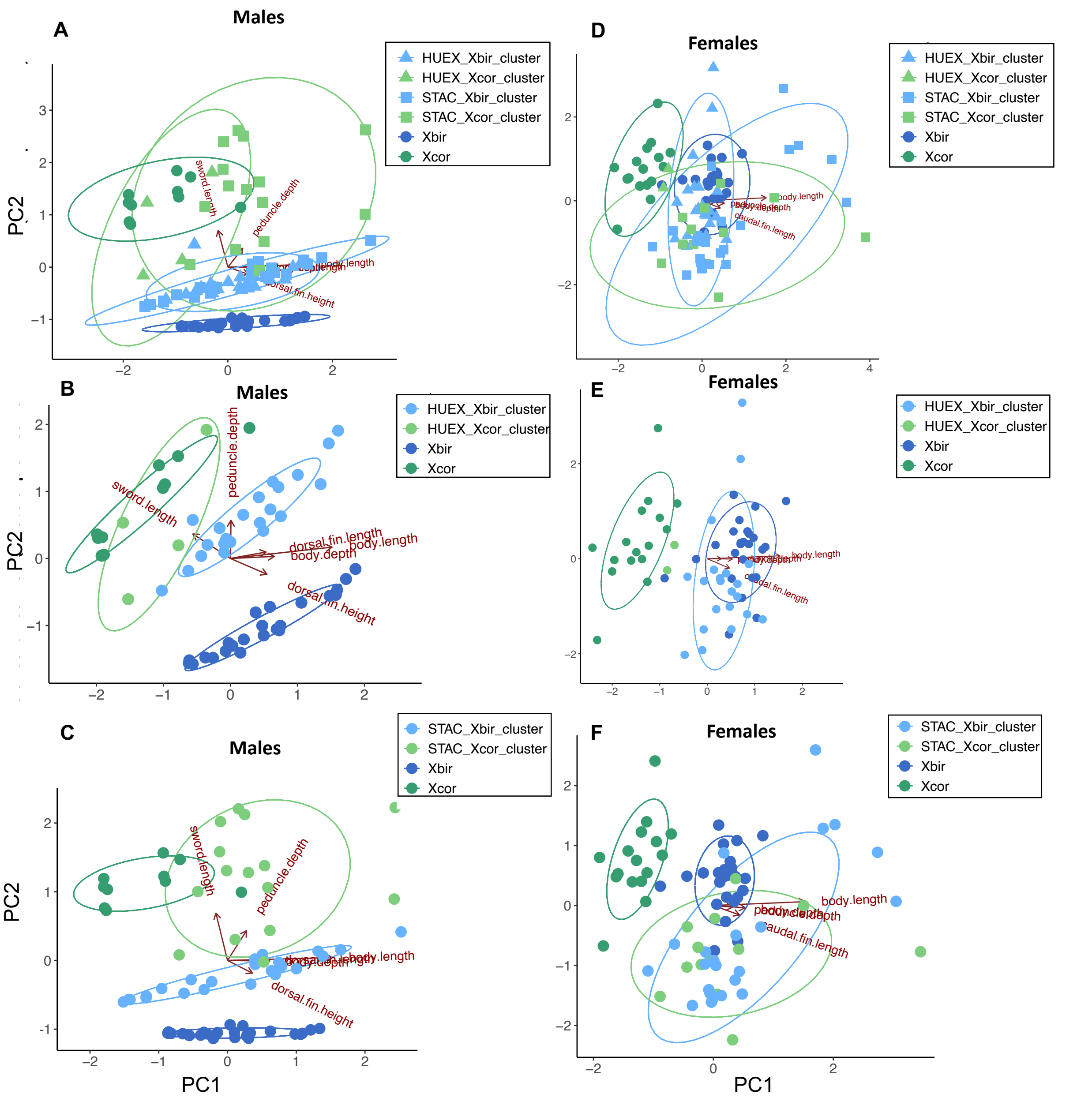


**Figure S5. Male and female PCAs with loadings plotted.** Principal Component plots of phenotypic data from all hybrid males (A), Huextetitla population males (B), Santa Cruz population males (C), all hybrid females (D), Huextetitla population females (E), and Santa Cruz population females (F) compared with parental species phenotypic data. Each point represents one individual and ellipses represent a 95% confidence interval.

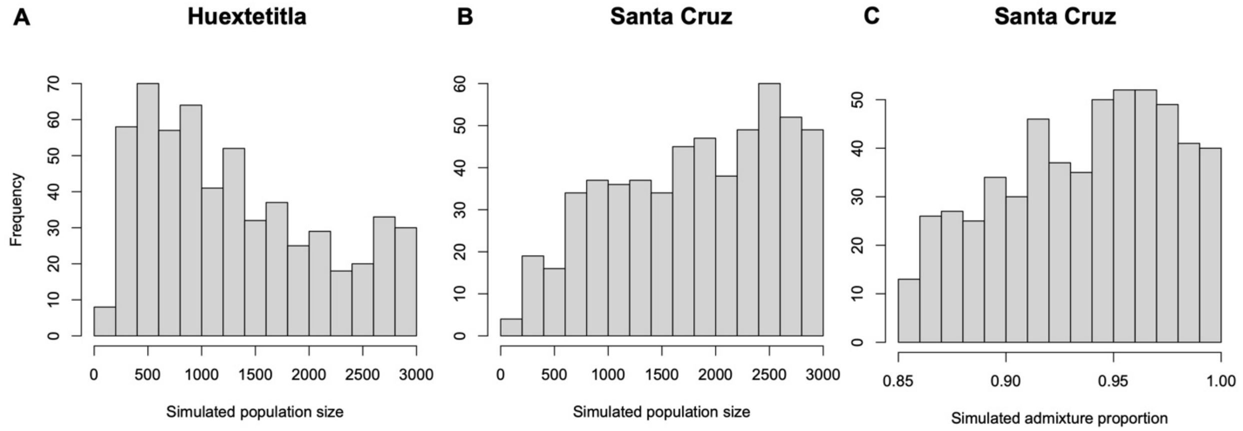


**Figure S6. Posterior distributions of parameters that were not well resolved in ABC simulations.** We used ABC simulations to infer the demographic history of the hybrid *cortezi* clusters at Huextetitla and Santa Cruz (see main text). Posterior distributions of hybrid population size for both Huextetitla (**A**) and Santa Cruz (**B**) were not well resolved but were skewed away from very small population sizes. **C**) Posterior distributions for simulated admixture proportion in Santa Cruz were also not well-resolved but are strongly correlated with accepted parameters for both time since initial admixture and migration rate (Fig. 3; Fig. S6).

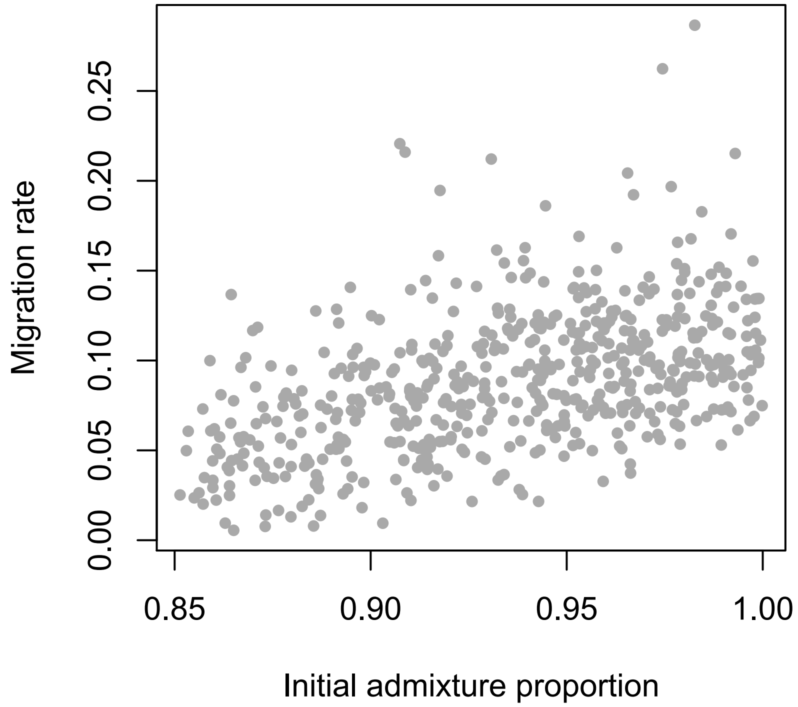


**Figure S7. Joint posterior distribution of migration rate and admixture proportion for ABC simulations fitting summary statistics for the Santa Cruz population.** In ABC simulations inferring the demographic history of the Santa Cruz hybrid population we observed strong correlations between accepted parameters for initial admixture proportion and migration rate (show here). We also observed similar correlations between accepted parameters for initial admixture proportion and time since admixture (Fig. 3).

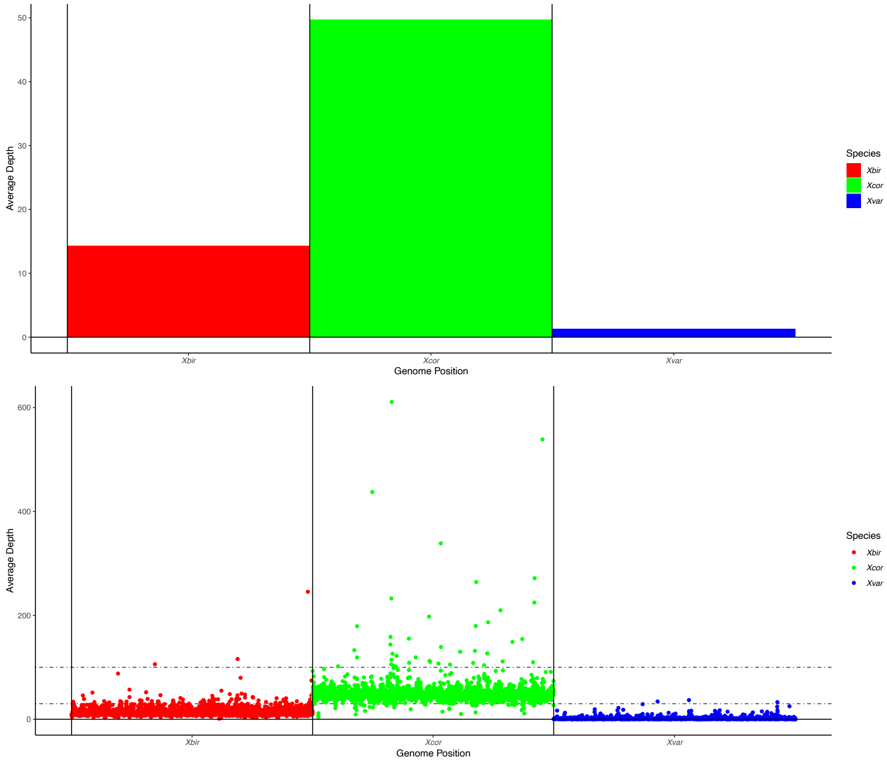


**Figure S8. SppIDer analysis results.** Mean depth of coverage for the *X. birchmanni* (Xbir - red), *X. cortezi* (Xcor - green), and *X. variatus* (Xvar) genomes when the deep-sequenced Huextetitla individual was competitively mapped to reference genomes of all three species (top). Coverage averaged in 200 kb windows (bottom) showed that a majority of the *X. cortezi* genome was covered at 30X-100X (dotted lines) while fewer of the *X. birchmanni* windows achieved this coverage, consistent with shorter ancestry tracts derived from *X. birchmanni*. Few windows in the *X. variatus* genome had above 0X coverage in this analysis.

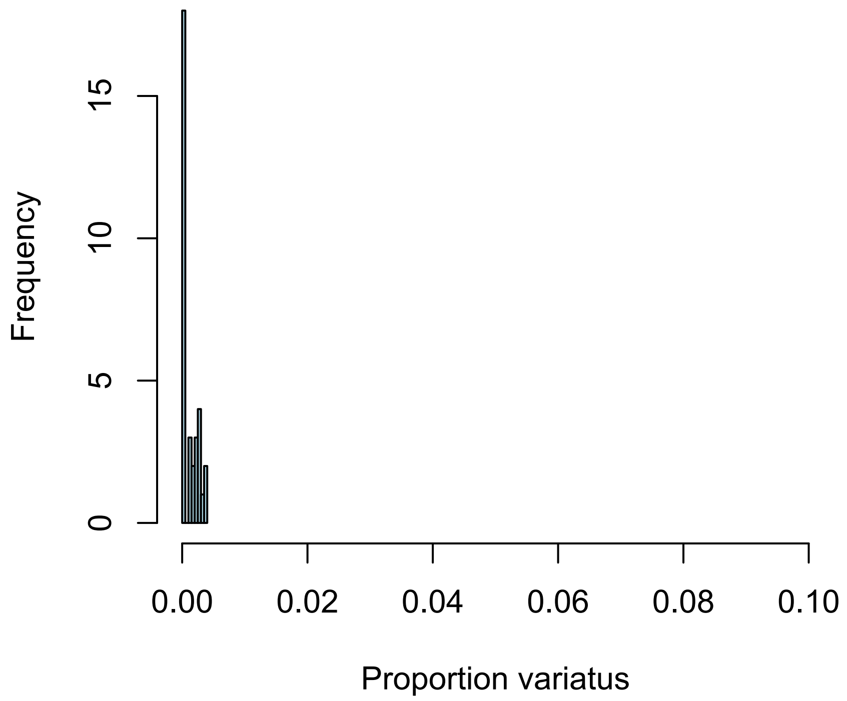


**Figure S9**. Histogram of inferred *X. variatus* ancestry genome-wide in a subset of individuals from the Santa Cruz and Huextetitla populations based on 3-way local ancestry calling.

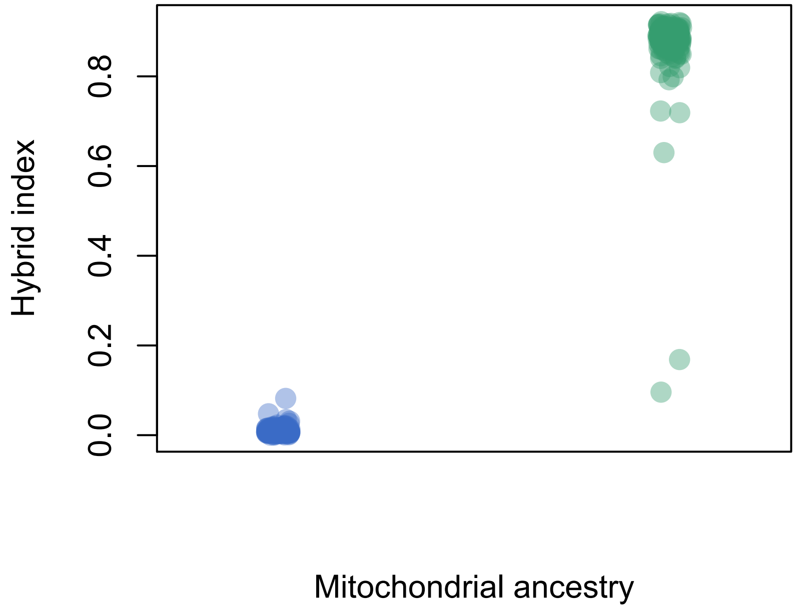


**Figure S10. Comparison between mitochondrial and nuclear ancestry in the Santa Cruz population.** Mitochondrial ancestry for each individual is plotted on the x-axis (blue – *X. birchmanni* mitochondria, green – *X. cortezi* mitochondria), relative to their genome-wide ancestry on the y-axis.

**Supporting Information References**


Corbett-Detig, R., & Nielsen, R. (2017). A Hidden Markov Model Approach for Simultaneously Estimating Local Ancestry and Admixture Time Using Next Generation Sequence Data in Samples of Arbitrary Ploidy. *PLOS Genetics*, *13*(1), e1006529. doi: 10.1371/journal.pgen.1006529

Cui, R., Schumer, M., Kruesi, K., Walter, R., Andolfatto, P., & Rosenthal, G. G. (2013). Phylogenomics reveals extensive reticulate evolution in *Xiphophorus* fishes. *Evolution*, *67*(8), 2166–2179. doi: 10.1111/evo.12099

Haynes, J. L. (1995). Standardized Classification of Poeciliid Development for Life-History Studies. *Copeia*, *1995*(1), 147. doi: 10.2307/1446809

Langdon, Q. K., Peris, D., Kyle, B., & Hittinger, C. T. (2018). sppIDer: A Species Identification Tool to Investigate Hybrid Genomes with High-Throughput Sequencing. *Molecular Biology and Evolution*, *35*(11), 2835–2849. doi: 10.1093/molbev/msy166

Li, H., & Durbin, R. (2009). Fast and accurate short read alignment with Burrows–Wheeler transform. *Bioinformatics*, *25*(14), 1754. doi: 10.1093/bioinformatics/btp324

Li, H., & Durbin, R. (2011). Inference of human population history from individual whole-genome sequences. *Nature*, *475*(7357), 493–496. doi: 10.1038/nature10231

McKenna, A., Hanna, M., Banks, E., Sivachenko, A., Cibulskis, K., Kernytsky, A., … DePristo, M. A. (2010). The Genome Analysis Toolkit: A MapReduce framework for analyzing next-generation DNA sequencing data. *Genome Research*, *20*(9), 1297–1303. doi: 10.1101/gr.107524.110

Powell, D. L., García-Olazábal, M., Keegan, M., Reilly, P., Du, K., Díaz-Loyo, A. P., … Schumer, M. (2020). Natural hybridization reveals incompatible alleles that cause melanoma in swordtail fish. *Science*, *368*(6492), 731–736. doi: 10.1126/science.aba5216

Schumer, M., Cui, R., Boussau, B., Walter, R., Rosenthal, G., & Andolfatto, P. (2013). An Evaluation of the Hybrid Speciation Hypothesis for Xiphophorus Clemenciae Based on Whole Genome Sequences. *Evolution*, *67*(4), 1155–1168. doi: 10.1111/evo.12009

Schumer, M., Cui, R., Powell, D. L., Rosenthal, G. G., & Andolfatto, P. (2016). Ancient hybridization and genomic stabilization in a swordtail fish. *Molecular Ecology*, *25*(11), 2661–2679. doi: 10.1111/mec.13602

Schumer, M., Powell, D. L., & Corbett-Detig, R. (2020). Versatile simulations of admixture and accurate local ancestry inference with mixnmatch and ancestryinfer. *Molecular Ecology Resources*, *20*(4), 1141–1151. doi: 10.1111/1755-0998.13175

Schumer, M., Xu, C., Powell, D. L., Durvasula, A., Skov, L., Holland, C., … Przeworski, M. (2018). Natural selection interacts with recombination to shape the evolution of hybrid genomes. *Science*, *360*(6389), 656–660. doi: 10.1126/science.aar3684
